# Piwi and piRNAs repress transcription of aberrant rRNA genes containing retrotransposon fragments

**DOI:** 10.1101/2025.04.30.651568

**Authors:** Elena A. Fefelova, Aleksei S. Shatskikh, Elena A. Mikhaleva, Yuri A. Abramov, Sergey A. Lavrov, Sergei A. Pirogov, Valentin A. Poltorachenko, Artem A. Ilin, Phillip D. Zamore, Mikhail S. Klenov

**Affiliations:** Institute of Molecular Genetics, Russian Academy of Sciences, Moscow 123182, Russia; Koltzov Institute of Developmental Biology, Russian Academy of Sciences, 26 Vavilov Street, Moscow 119334, Russia; Department of Molecular Biosciences, Stockholm University, The Wenner-Gren Institute, Stockholm, Sweden; Department of Microbiology, Tumor and Cell Biology, Karolinska Institutet, Stockholm, Sweden; RNA Therapeutics Institute, University of Massachusetts Chan Medical School, 368 Plantation Street, Worcester, MA 01605, USA; Howard Hughes Medical Institute, University of Massachusetts Chan Medical School, 368 Plantation Street, Worcester, MA 01605, USA

**Keywords:** rDNA, ribosome, transposon, piRNA, Piwi, rRNA, nucleolus, regulation of transcription, gene silencing

## Abstract

The genomes of eukaryotes comprise clusters of repeated rRNA genes, including defective copies that are silenced by poorly understood mechanisms. In *Drosophila melanogaster*, many 28S rDNA genes contain insertions of R1 or R2 retrotransposons. R2 elements can only transcribe as part of the pre-rRNA and then excise by the R2 ribozyme. Some rRNA genes carry truncated insertions, creating defective rRNA from which the R2 sequence cannot be excised. Here, we report that in *Drosophila* ovaries, the nuclear protein Piwi loaded with PIWI-interacting RNA (piRNA) is required to repress transcription of rDNA units with extensively truncated R2 insertions (R2short). Without Piwi, R2short-rDNA generates stable aberrant 28S rRNA containing ∼200 nt of R2 sequence. These rRNAs are excluded from cytoplasmic ribosomes and instead accumulate within the nucleoli of germ cells, where they may disrupt nucleolar homeostasis. Overall, our results show the involvement of the piRNA pathway in quality control of ribosomal transcription.

## Introduction

In eukaryotes, rDNA clusters, also known as nucleolus organizing regions (NORs), comprise tandemly repeated rRNA gene units that are transcribed by RNA polymerase I (Pol I) within the nucleolus, producing the most abundant transcripts in the cell.^1^ Each unit generates a precursor transcript (pre-rRNA) in which the mature 18S, 5.8S, and 28S rRNAs are separated by internal transcribed spacers (ITSs). Typically, only a fraction of rDNA units are transcriptionally active. Both entire NORs and individual units within a NOR can be subjected to transcriptional repression.^2–4^ Selective silencing of specific rDNA copies is thought to regulate rRNA synthesis in response to cellular conditions and to repress defective rRNA genes. How transcription of individual rDNA units is repressed remains poorly understood.

rRNA genes are often damaged by transposable element (TE) insertions.^5^ In particular, rDNA units in the genomes of various animals are interrupted by the non-LTR retrotransposons R1 or R2^6^. In *Drosophila melanogaster*, from 30% to 80% of rDNA units contain insertions of these elements depending on the line.^7^ R2 transposons integrate exclusively into specific site within the 28S rDNA due to the unique properties of the R2-encoded protein (Fig. 1A). R1 elements also have a preferred integration site within the 28S rDNA, located ∼70 bp downstream of the R2 insertion site, but they are also found in other genomic regions outside rDNA units.^7,8^

**Fig. 1.**
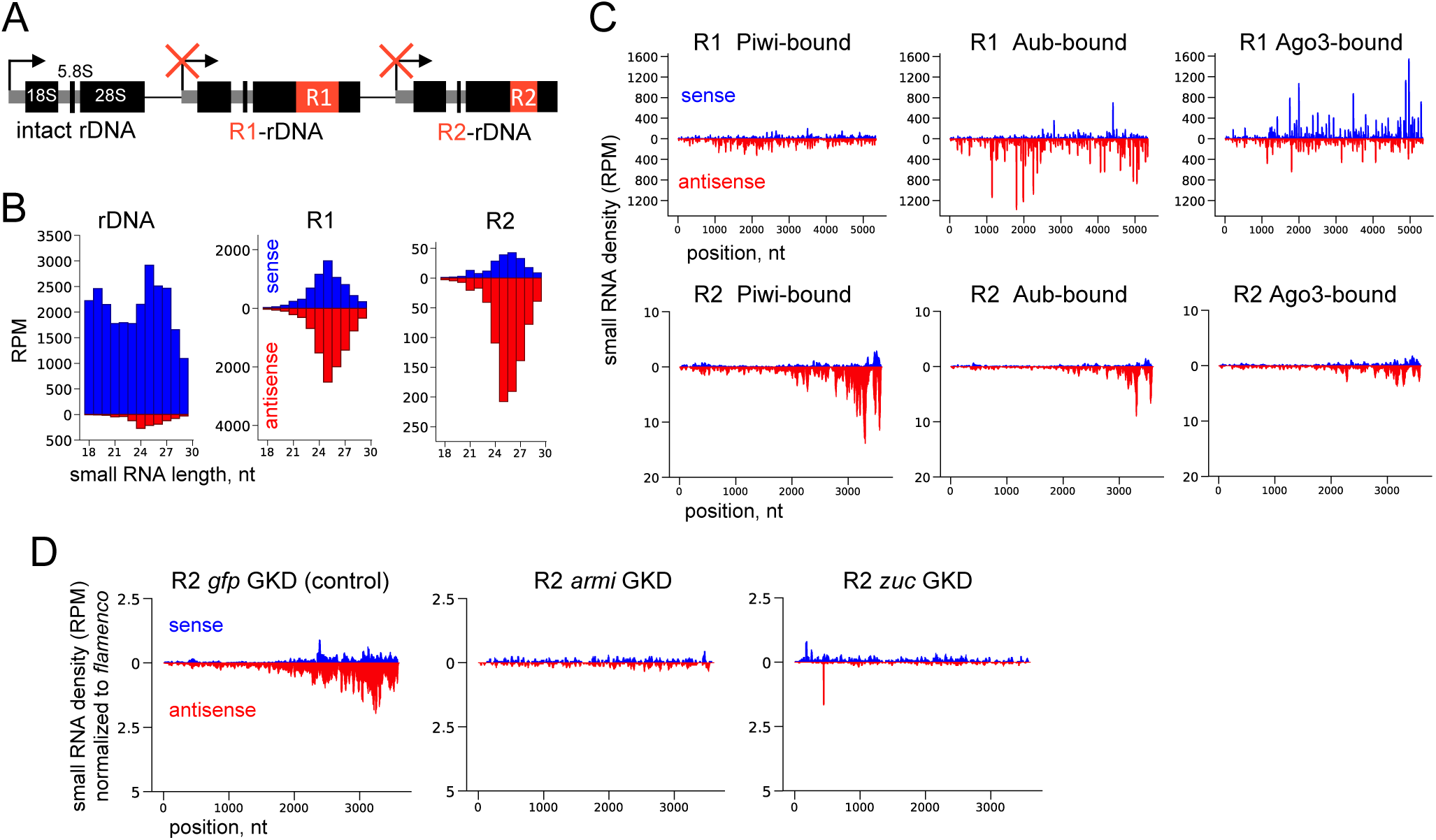
Germline Piwi is loaded with piRNAs antisense to R2 elements. **A.** Schematic of rDNA units in *D. melanogaster*. **B.** Size distribution of total ovarian small RNAs mapped to rDNA and R1 and R2 elements in the *w*^1118^ control line. **C.** Coverage of R1 and R2 elements by small RNAs associated with immunoprecipitated Piwi, Aub, and Ago-3 proteins. **D.** Coverage of R1 and R2 by small RNAs in MTD-GAL4-driven germline knockdowns (GKD) of GFP (control) and two piRNA biogenesis factors: Armi and Zuc normalized to piRNAs derived from the somatic *flamenco* piRNA cluster.

R2 elements lack promoters and can only be transcribed as a part of the pre-rRNA from which they auto-excise via a self-cleaving ribozyme near the R2 5′ end.^9^ In *D. melanogaster*, rDNA loci reside on the X and Y chromosomes, typically containing 100–300 and 30–250 rDNA units, respectively.^10^ rDNA copy number may be reduced due to sporadic recombination.^11,12^ R2 element expression can benefit cells with low rDNA copy number, because DNA breaks created by the R2 endonuclease can initiate recombinational recovery of rDNA copy number in germline stem cells.^13,14^ Nonetheless, R2 acts predominately as a selfish genetic element, damaging a large fraction of rRNA genes.^6^ In addition to transposition-competent, full-length R1 (5.3 kb) and R2 (3.6 kb), many rDNA units carry truncated insertions resulting from incomplete reverse transcription of retrotransposon RNA.^15–17^

The expression of rDNA units with R1 and R2 insertions (hereafter, R1-rDNA and R2-rDNA) varies among *Drosophila* strains^15,18,19^ but is typically orders of magnitude lower than that of intact rRNA genes.^20–23^ The absence of the nucleolar protein Nopp140^24^ and, paradoxically, impairment of the Pol I-specific transcription initiation complex^22^ increase transcription of R2-rDNA, likely as a compensatory response to reduced synthesis of functional rRNA. Consistently, R2-rDNA expression is also elevated in cells with low rDNA copy number.^14,25,26^ A slight derepression of R1 and R2 occurs upon the loss of certain chromatin-associated proteins, including histone H1,^27^ lamin,^28^ CTCF,^29^ H3K9 methyltransferases, and HP1^22,30^. In contrast, loss of SUMO (small ubiquitin-like modifier) results in much stronger derepression of R1 and R2 than other transposable elements in *Drosophila*.^30,31^ Despite these observations, the molecular mechanisms by which flies recognize and selectively silence rDNA units with transposon insertions remain unknown.

In animals, including flies, the PIWI-interacting RNA (piRNA) pathway silences TEs in the germline.^32,33^ 18–35-nt piRNAs are produced from single-stranded transcripts and bind members of the PIWI sub-family of Argonaute proteins. In *D. melanogaster*, the cytoplasmic PIWI proteins Aubergine (Aub) and Argonaute3 (Ago3) both amplify the abundance of pre-existing piRNAs and cleave complementary target RNAs.^34–36^ By contrast, Piwi, the third *Drosophila* PIWI protein, mediates piRNA-guided transcriptional silencing in the nucleus by recruiting a Histone H3 lysine 9 trimethylase and other factors that generate heterochromatin.^37–43^ The piRNA pathway in *D. melanogaster* is active primarily in gonads to prevent TE transpositions in the germline genome. Along with TE silencing, ovarian germ cells must also meet the high demand for ribosome production required for oocyte development and the early phases of embryogenesis.^44^ Piwi is localized throughout the cell nucleus, including the nucleolus of germ nurse cells, suggesting a potential role in the repression of interrupted rRNA genes in the germline.^45^ On the other hand, rough estimates based on RNA-seq suggest that R1 and R2 expression is typically not significantly altered in the ovaries of piRNA pathway mutant flies.^31,43,46,47^

Here, we report that Piwi silences transcription of rDNA units containing highly truncated ∼200 bp fragments of R2 element (R2short). We found that piRNAs act also on rDNA interrupted by longer or full-length R1 or R2 insertions, but this repression is redundant with an as yet unidentified mechanism that silences rDNA units with insertions longer than 200 bp. We show that in *piwi* mutants, R2short-rDNA units produce aberrant 28S rRNA with embedded R2 sequence; these transcripts accumulate in pre-ribosomes within nucleoli but do not appear in the cytoplasmic pool of ribosomes. Our results expand the known functions of Piwi-mediated transcriptional silencing to include a subclass of defective rRNA genes, thereby contributing to nuclear RNA homeostasis.

## Results

### In ovarian germ cells, nuclear Piwi is loaded with piRNAs antisense to R2 elements

Reanalysis of published high-throughput small RNA sequencing data for fly ovaries revealed small RNAs mapping to R1 and R2 retrotransposons. As noted previously,^35^ we detected piRNA-sized (23–30 nt), as well as less abundant siRNA-sized (21 nt) RNAs, matching or antisense to the R1 or R2 consensus sequences (Fig. 1B, S1A, and S1B). Aub and Piwi typically bind piRNAs with a 5′ uridine, and more than half of the R1 and R2 small RNA sequences begin with U, supporting their identification as piRNAs (Fig. S1C). Consistent with this, oxidized small RNA-seq data show that most R1 and R2 small RNAs have blocked 3′ ends (Fig. S1D), a hallmark of piRNAs.^35^ Notably, R2 piRNAs exhibit a strong bias for the antisense orientation in all eight lines analyzed (Fig. 1B, S1A, and S1B, Table S1), suggesting their role in targeting pre-rRNA-derived sense R2 transcripts.

Analysis of data sets for small RNA coimmunoprecipitating with each of the three fly PIWI proteins^48–50^ revealed that R2 piRNAs are bound predominantly to Piwi, whereas R1 piRNAs were bound mainly to Aub in some data sets and Piwi in others (Fig. 1C and S2, Table S1). Interestingly, R2 piRNAs complementary to sequences at the 3′ end of the TE predominated among both total and Piwi-bound small RNA (Fig. 1C, S1B, and S2C). Mapping of small RNAs to rDNA sequence revealed that PIWI proteins are loaded with highly abundant RNAs derived from mature rRNAs (Fig. S2A and S2C, Table S1). However, only a small fraction of piRNAs were complementary to rRNA sequences. Unlike sense rRNA-derived piRNAs, antisense piRNAs span the entire rRNA gene, covering mature rRNAs as well as external (ETS) and internal transcribed spacer (ITS) sequences (Fig. S1C and S2C).

piRNA biogenesis relies on two mechanisms: ping-pong amplification and the phased piRNA pathway, in which the Zuc nuclease generates tail-to-head strings of piRNAs loaded mostly into Piwi.^49,51^ In flies, phased piRNA production requires the putative RNA helicase Armitage (Armi), but piRNA ping-pong amplification persists in *armi* mutants.^36,52^ Compared to control *armi^Delta^*^1^/+ and *w*^1118^ flies, R2 piRNAs were ∼30- and ∼100-fold reduced (p < 0.001) in datasets from *armi^72.1^/armi^Delta^*^1^ and *w*^1118^*; armi^72.1^/armi^G728E^*mutant ovaries, respectively (Fig. S3A and S3B). In contrast, piRNAs derived from R1 were only decreased ∼2-fold in *armi^72.1^/armi^G728E^* ovaries (p < 0.001) (Fig. S3B). Aub, Ago3, and ping-pong amplification are germline-specific in flies, but an abbreviated piRNA pathway dependent on Piwi silences *gypsy* family TEs in somatic follicle cells.^35,36^ R2 piRNAs were reduced >10-fold in germline-specific knockdowns of *zuc* and *armi*, when normalized to piRNAs derived from the somatic *flamenco* piRNA cluster (Fig. 1D and S3C). Thus, R2 piRNAs are mainly produced from the phased piRNA pathway in germ cells.

### Piwi mutations lead to derepression of rRNA genes with extensively truncated R2 insertions

R2-rDNA units show considerable structural variability among fly lines. Like other non-LTR retrotransposons, R2 elements can be truncated from the 5′ end due to incomplete reverse transcription. Consequently, some rDNA units contain R2 insertions that vary in the extent of 5′ truncation but retain identical 3′-end sequences (Fig. 2A).^15,17,53^ To investigate Piwi role in R2-rDNA silencing, we first examined the presence of truncated R2 insertions in various *piwi* loss-of-function mutants. Long-extension PCR of genomic DNA, using primers flanking R1 and R2 insertion sites in 28S rDNA (Fig. 2B),^17^ followed by sequencing of individual PCR products, revealed the highly truncated ∼200 bp R2 fragments in the *piwi^2^* strain^54^ and in strains carrying the *piwi^2^-*derived X chromosome rDNA cluster (Fig. 2B and S4). Hereafter, we designate these insertions as R2short and the corresponding rDNA units as R2short-rDNAs. Other analyzed *piwi* loss-of-function lines, such as *piwi^Nt^* allele,^38^ lacked R2 insertions shorter than 1 kb (Fig. 2B and S4). Of note, some *D. melanogaster* lines were previously shown to contain similar sized R2short isoforms, including 167-bp^15^ and 180-bp^22^ variants. By nanopore sequencing we found that the *piwi^2^* X-chromosome rDNA cluster contains approximately 200 rDNA units, of which 25–30% are intact, 50–55% harbor R1 insertions and 20% have R2 insertions, including ∼10 full-length (3.6 kb) and ∼30 truncated R2 isoforms ranging in size from ∼0.2 to ∼3.1 kb. Among them, four rDNA units contain ∼200 bp insertions, designated R2short #1–4 (Fig. 2C and S5). Their lengths and sequences, including a non-templated 10–12 nt poly-T tract, match the R2short variants identified by PCR (Fig. S5C). The four R2short-rDNA units are distributed at different locations within the rDNA cluster (Fig. 2C, S5A, and S5B). Two of them (#1 and #2) are separated by a 130-kb region that includes six rDNA units with R1 insertions and one with a 540-bp R2 insertion (Fig. 2C). Within the accuracy of nanopore sequencing, all four R2short-rDNAs appeared to be full-length rRNA genes displaying no other abnormalities in either their pre-rRNA-producing regions or upstream promoter elements (Fig. 2C, S5A, and S5B). Individual R2short insertion sequences were 94–97% identical (Fig. S5C), and we cannot exclude that sequence differences reflect nanopore sequencing errors. It is unlikely that multiple nearly identical R2short insertions—produced by incomplete reverse transcription during transposition—occurred independently in the same line, suggesting that they arose during rDNA copy number expansion.

**Fig. 2.**
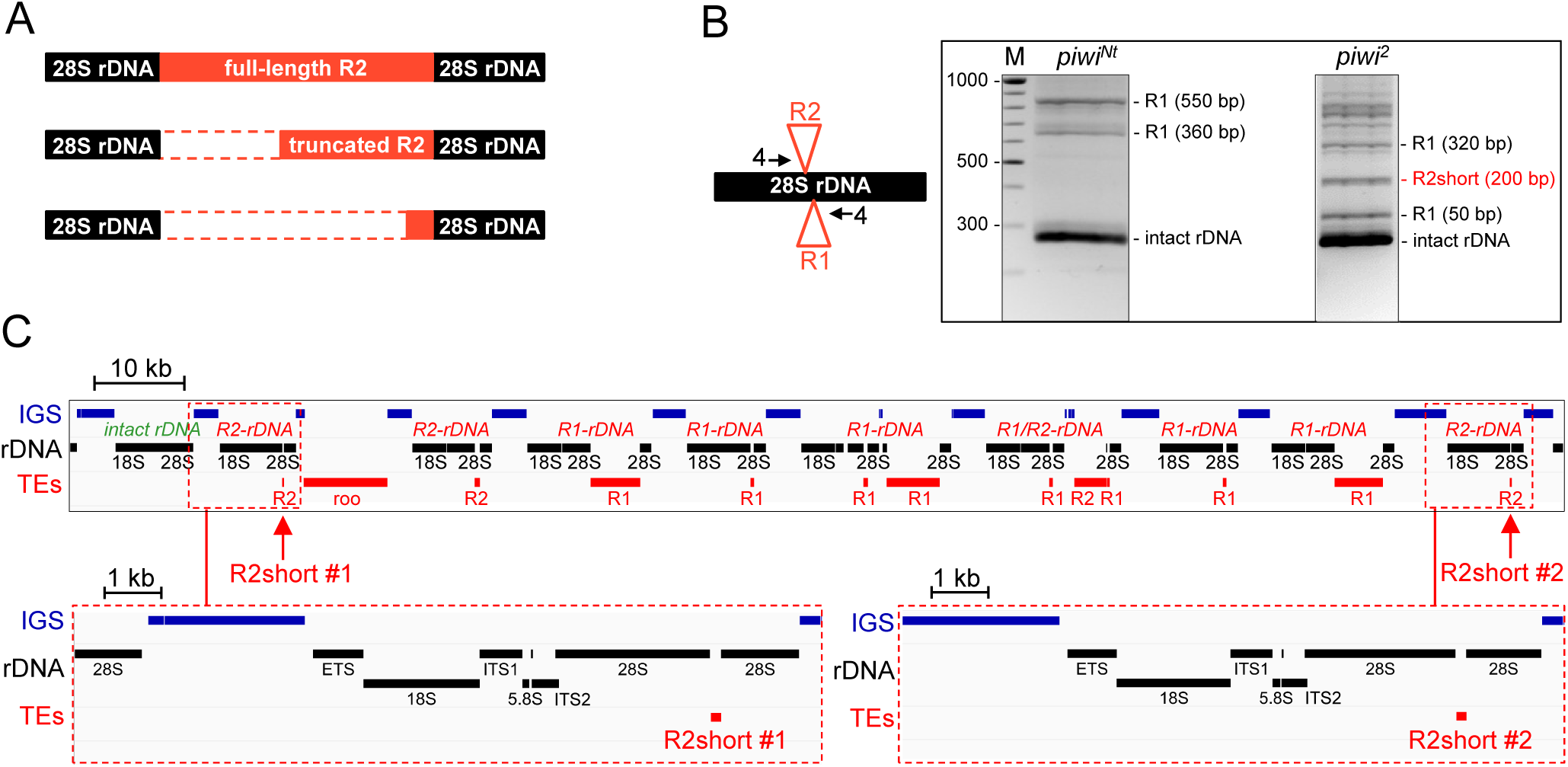
Characterization of rDNA units containing R2short insertions. **A.** Schematic of truncated R2 insertions within rDNA. **B.** Left: Location of primers #4 (black arrows) for long-extension PCR analysis of R1 and R2 insertions. R1 elements typically integrate 74 bp downstream of the R2 insertion site. Right: gel image showing the most highly truncated R1 and R2 insertions in the genomic DNA of *piwi^Nt^* and *piwi^2^* females. The intact 28S rDNA, lacking insertions, is detected as 268 bp PCR product (lower band). For rDNA units containing insertions, the PCR product size increases by the insertion length. M: 1kb DNA ladder. **C.** Top: IGV browser screenshot showing a 180-kb fragment of the rDNA cluster in the *piwi^2^* line, obtained by juxtaposition of two uniquely overlapped nanopore reads. Positions of intergenic spacers (IGS, blue), rDNA (black), and TE insertions (red), including R1 and R2, are indicated. Bottom: enlarged view of rDNA-R2short #1 and R2short #2 units.

By RT-qPCR we assessed changes in R1 and R2 expression in different *piwi* loss-of-function mutants compared to their heterozygous sisters, as well as in *piwi* RNAi germline knockdown (Fig. 3A, 3B, and S6A). Intriguingly, we observed >20-fold derepression for R2 3′-terminal regions in mutants with R2short insertions, but not in lines without R2short (Fig. 3A). Levels of RNAs derived from regions located further upstream in the R2 sequence showed ≤4-fold increase in all *piwi* mutants (Fig. 3A).

**Fig. 3.**
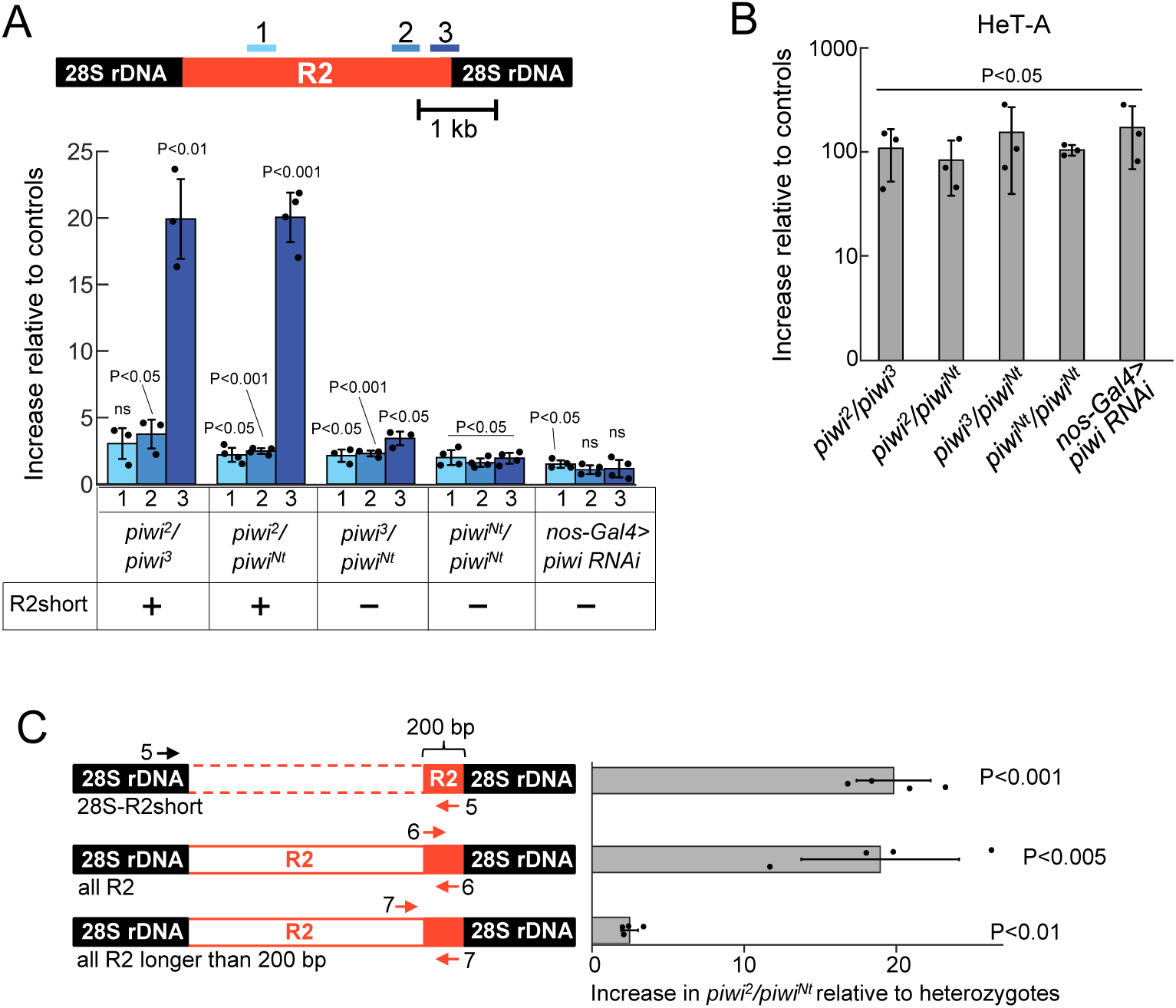
Piwi mutations derepress R2short-rDNA units. **A.** Increase (fold change) in RNA levels at different regions of the R2 sequence measured by RT-qPCR in *piwi* mutant combinations compared to heterozygous sisters carrying the same X chromosomes. Numbers indicate primer pairs. The presence (+) or absence (-) of R2short insertions is indicated based on genomic PCR analysis. **B.** Increase of HeT-A retrotransposon RNA level, used as a positive control of transposon derepression in analyzed *piwi* mutants. **C** Increase in RNA levels measured by RT-qPCR in *piwi^2^/piwi^Nt^* mutant compared to heterozygous sisters (mix of *piwi^2^/+*; *piwi^Nt^/+*). Numbers and positions of primers are shown. Data in all panels are normalized to *rp49* mRNA. p-values are from a two-tailed t-test; ns = not significant.

These results suggest that Piwi absence more strongly derepresses R2short compared to other R2 isoforms. Since R2short insertions lack the ribozyme required to excise transposon RNA from the upstream rRNA, expression of R2short-rDNA can be specifically detected using a forward primer to the upstream 28S rRNA sequence and a reverse primer to the R2 3′-terminal sequence (Fig. 3C, S6B, and S6C). In *piwi^2^/piwi^Nt^* ovaries, the 28S-R2short junction RNA increased by ∼20-fold ± 3 (mean ± SD) compared to heterozygous siblings (p < 0.001) (Fig. 3C). Similar increase was observed with primers located within 200 bp of the 3′ end of R2, which detects expression of all R2 variants including R2short. Overall abundance of RNA from all R2 variants longer than 200 nt increased ∼2-fold in *piwi^2^/piwi^Nt^* ovaries (Fig. 3C). Of note, the transheterozygous *piwi^2^/piwi^Nt^* genotype combines a *piwi* null-mutation with the *piwi^Nt^* allele, in which Piwi lacks a nuclear import sequence. *piwi^2^/piwi^Nt^* flies are defective in transcriptional piRNA-directed silencing, because Piwi^Nt^-piRNA complexes cannot be imported into the nucleus.^38^ Despite ∼20-fold R2short derepression, R2 piRNA abundance was reduced by only ∼1.4-fold in *piwi^Nt^/piwi*^2^ ovaries (Fig. S3E and Table S1). This indicates that cytoplasmic production of R2 piRNAs remains relatively unaffected, whereas silencing of R2short-rDNA depends on Piwi nuclear localization.

Why do *piwi* mutations derepress R2short-rDNA but not long R2? Perhaps Piwi-piRNAs can target rDNA units with R2 insertions of any length, but long R2-rDNAs remain silenced by alternative pathways independent of Piwi. To test the idea that long insertions can be targeted by Piwi, we took advantage of the increased expression of R2 elements in germ cells depleted of Udd,^22^ a component of the SL1-like rDNA transcription initiation complex^55^. We measured R1 and R2 transcript levels in flies in which Piwi, Udd, or both Piwi and Udd were depleted by RNAi; all genotypes contained the same X chromosomes. Long R2 transcripts increased ∼13-fold ± 9 (mean ± SD) when Udd was depleted (Fig. S6D). When both Piwi and Udd were depleted, long R2 RNA increased ∼80-fold ± 10. Thus, when long R2 expression is increased as Udd is compromised, Piwi still constrains the extent of that increase. Similarly, R1 RNA increased 3-fold upon germline knockdown of Udd, but ∼30-fold when both Udd and Piwi were depleted (Fig. S6D). These results suggest that the piRNA pathway acts as a backup mechanism constraining the expression of long R2- and R1-rDNAs. However, Piwi is insufficient to fully silence R2-rDNAs when they are robustly activated.

### R2short-rDNA units produce aberrant 28S rRNAs with embedded R2short sequence

In full-length R2 elements, the R2 ribozyme detaches R2 from the upstream 28S rRNA sequence.^9^ In contrast, the 3′ R2 sequence lacks recognizable ribozyme structures, and the mechanism of R2 RNA 3′ end formation is unknown.^6^ We found that transcription of R2short-rDNA proceeds to the downstream rRNA generating an R2 fragment sandwiched between two 28S rRNA parts. First, the abundance of transcripts in which the R2 3′ end was joined to the downstream 28S rRNA sequence (3′R2-28S transcripts) was 100–1000-fold higher in *Drosophila* lines containing R2short compared to lines without it, whereas the levels of R2 body RNA (i.e., sequences 5′ to the 200 bp R2 3′- terminal sequence) were similar in all lines tested (Fig. 4A), suggesting that R2short- rDNAs are the major source of 3′R2-28S transcripts. Second, primer-specific reverse transcription followed by qPCR in *piwi^2^*/*piwi^Nt^* ovaries revealed that 3′R2-28S transcripts are generated by R2short-rDNA units ∼60-fold more frequently than by longer R2 insertions (Fig. S7). Third, the R2 junctions with the downstream rRNA were present at levels similar to R2short junctions with the upstream rRNA (28S-R2short) (Fig. 4A).

**Fig. 4.**
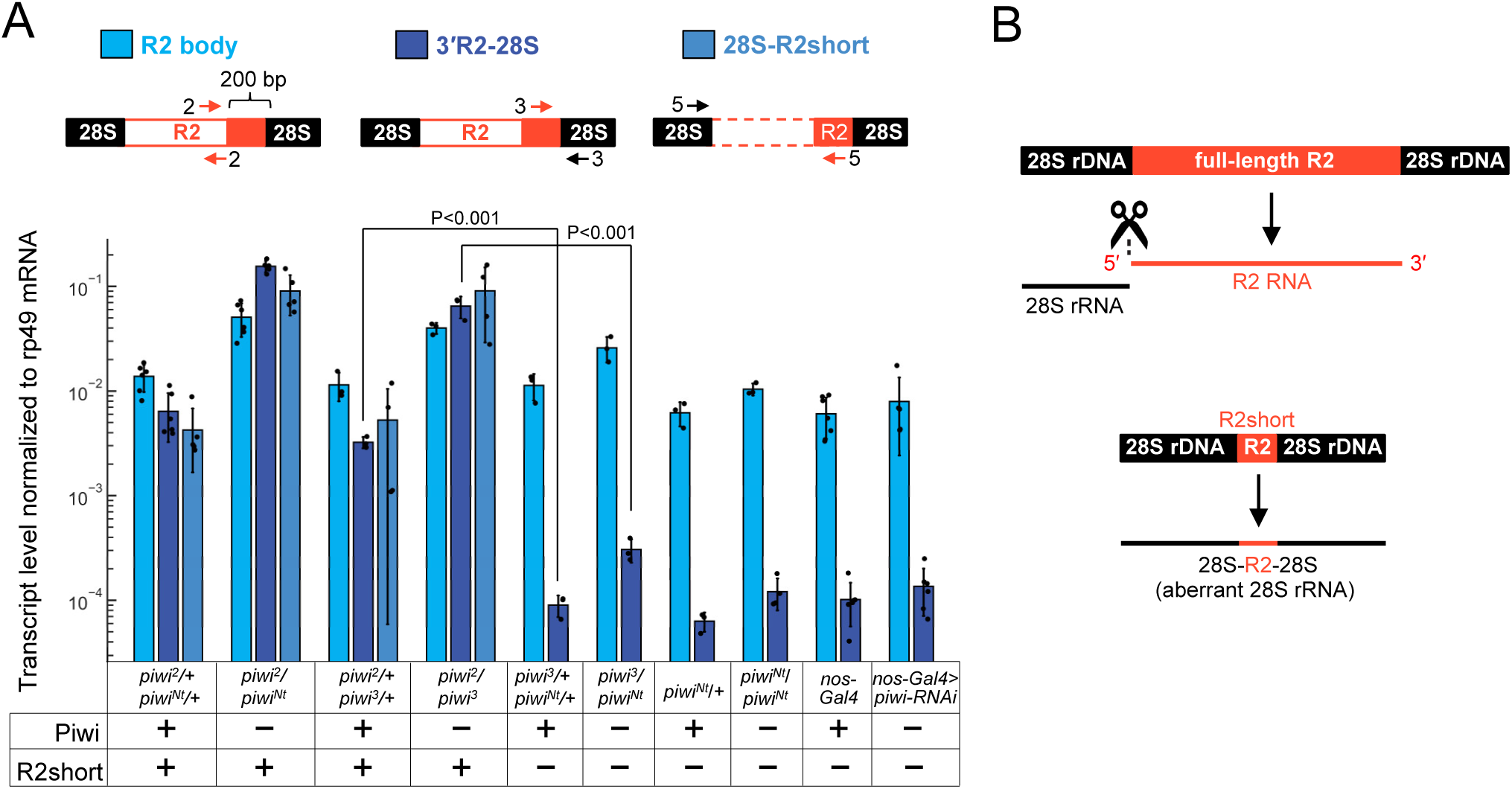
R2short-rDNA units produce aberrant 28S rRNAs with embedded R2short sequence. **A** The abundance of R2 body RNA and 3′R2-28S transcripts, containing junctions of the R2 3′ sequence and downstream 28S rRNA, measured by RT-qPCR in ovaries of different *piwi* mutants (Piwi -) and their heterozygous sisters (Piwi +). The presence (+) or absence (-) of R2short insertions is indicated based on genomic PCR analysis. Numbers and positions of primers are shown. Data are normalized to *rp49* mRNA. p-values are from a two-tailed t-test. **B** Cartoon depicting the release of full-length R2 transcripts from 28S rRNA, alongside the lack of release for R2short elements (not to scale).

Given that R2short cannot be excised from upstream rRNA, we concluded that most R2short transcripts remain trapped in rRNA on both sides (Fig. 4B). Thus, *piwi* mutation leads to derepression of the aberrant 28S rRNA containing a non-excisable 200-nt R2 insertion.

### Piwi prevents accumulation of R2short-rRNA in germ cell nucleoli

The R2 insertion is located within a conserved sequence in domain IV of the large ribosomal subunit.^56^ Do aberrant 28S rRNAs containing R2short sequence accumulate in mature ribosomes or do nuclear quality control mechanisms prevent the assembly or export of such defective complexes? We purified nuclei and cytoplasm from *piwi^2^/piwi^Nt^* ovaries and then separated cytoplasmic ribosomes from cytosol by ultracentrifugation.

The pellet contained cytoplasmic ribosomes, including both polysomes and monosomes, whereas the post-ribosomal cytoplasmic supernatant corresponded to cytosol. For each subcellular fraction, we evaluated the relative abundance of various rDNA-derived transcripts by RT-qPCR using equal amounts of total RNA. As expected, the relative abundance of intact 28S rRNA devoid of TE insertions was ∼1000 times higher in the cytoplasmic ribosomes than in the cytosol, and pre-rRNA was >100 times more abundant in nuclei compared to cytoplasmic ribosomes or cytosol (Fig. 5A).

**Fig. 5.**
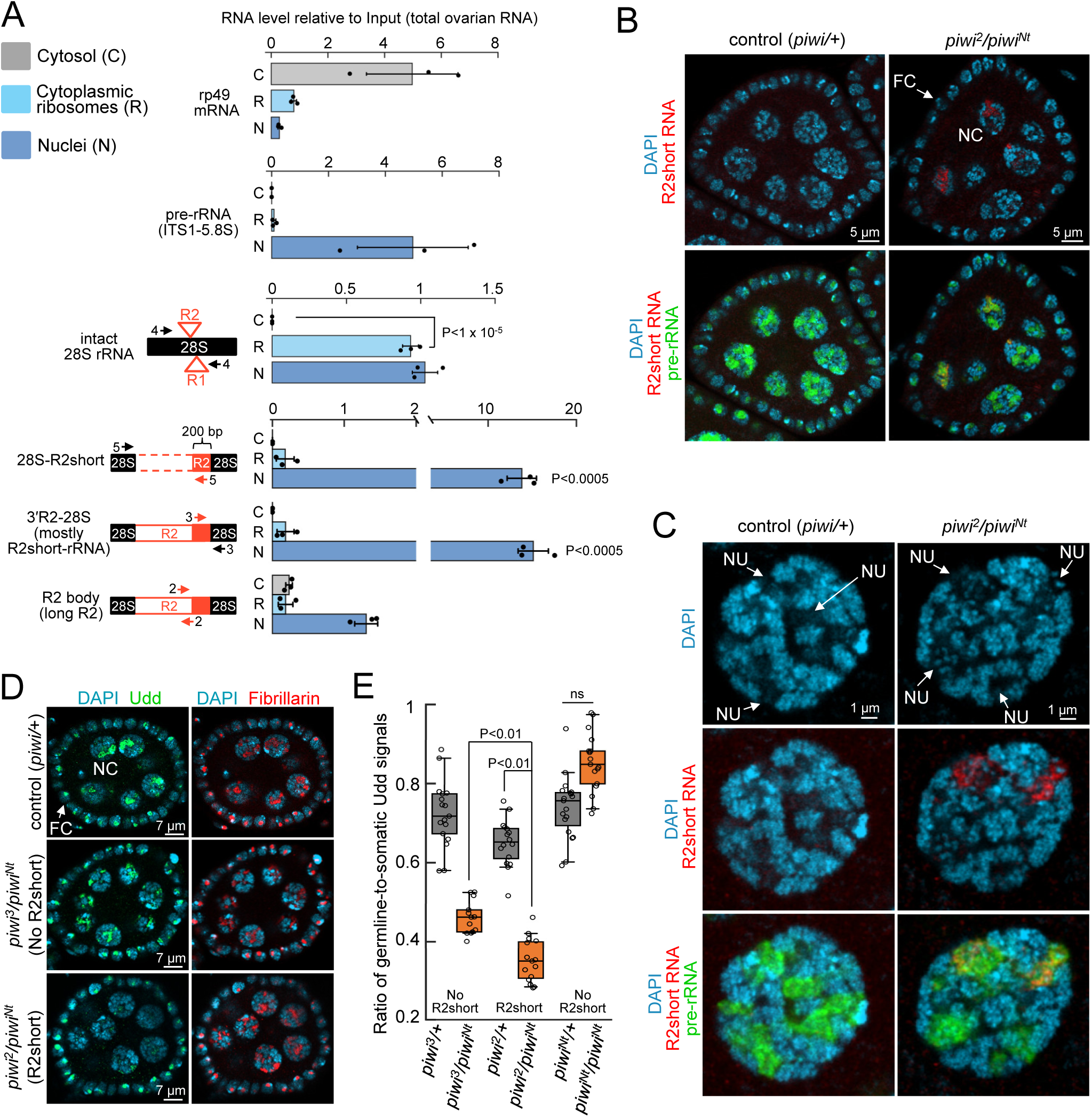
R2short-rRNA accumulates in germ cell nucleoli. **A** Relative abundance of transcripts in cytosol (C), cytoplasmic ribosomes (R), and nuclei (N) normalized to total ovarian RNA (Input) in *piwi^2^/piwi^Nt^* ovaries. The level of R2short-rRNA was assessed by measuring the amounts of both 28S-R2short and 3’R2-28S junctions. Numbers and positions of primers are shown. p-values are from a two-tailed t-test. **B** R2short RNA and pre-rRNA FISH in stage 4 egg chambers of *piwi^2^*/*piwi^Nt^* mutants and control heterozygotes. Germline nurse cells (NC) and somatic follicle cells (FC) are indicated. **С** Airyscan 2 super-resolution images of R2short RNA and pre-rRNA FISH within single nurse cell nuclei. Nucleolar regions are designated as NU. **D** Immunostaining for the nucleolar markers fibrillarin and Udd in stage 4 egg chambers of *piwi* mutants, either containing or lacking R2short-rDNA. Udd shows impaired localization in nurse cells (NC) of *piwi* mutants containing R2short-rDNA, but not in follicle cells (FC). **E** Quantification of the germline-to-somatic Udd signal ratio in control and mutant ovaries: *piwi^3^/piwi^Nt^* (lacking R2short-rDNA); *piwi^2^/piwi^Nt^* (containing R2short-rDNA); *piwi^Nt^/piwi^Nt^* (lacking R2short-rDNA). Each data point represents a single egg chamber of stage 4-5. Whiskers represent the 5th to 95th percentiles. p-values are from two-tailed Mann-Whitney U test; ns = not significant.

R2short-rRNA measured at both 28S-R2short and 3′R2-28S junctions was found predominantly in nuclei (Fig. 5A). Little R2short-rRNA was detected in the cytoplasmic ribosome fraction (∼2% ± 2) or in the cytosol (0.004% ± 0.004) (Table S2). Long R2 RNA was also mostly nuclear. However, the relative abundance of long R2 RNA in the cytosol was >100-fold higher than for R2short-rRNA, indicating that some long R2 RNA is exported from the nucleus (Fig. 5A).

Assembly of rRNA with ribosomal proteins and maturation of ribosomal subparticles occurs within the nucleolus.^57^ Purification of pre-ribosomes from nuclear lysates showed that approximately 80% ± 20 of R2short-rRNA was found in the pre-ribosomal pellet, similar to the proportion of nuclear intact 28S rRNA (Table 1). Thus, R2short-rRNAs are incorporated into maturating ribosomes, but these are either not exported from the nucleus or undergo rapid degradation in the cytoplasm. By contrast, ∼90% of nuclear long R2 transcripts were found in the nuclear supernatant after ultracentrifugation (Table 1), indicating they are not associated with pre-ribosomes.

**Table 1.** The percentage of molecules in nuclear pre-ribosomes and nuclear supernatant as fraction of cleared nuclear lysate.

|  | Fraction of cleared nuclear lysate (%) |  |
| --- | --- | --- |
|  | Sediment (pre-ribosomes) | Supernatant |
| rp49 mRNA | 2 ± 2 | 98 ± 2 |
| intact 28S rRNA | 70 ± 30 | 30 ± 20 |
| 28S-R2short | 80 ± 20 | 20 ± 20 |
| 3'R2-28S (mostly R2short-rRNA) | 80 ± 20 | 20 ± 20 |
| long R2 (R2 body2) | 10 ± 10 | 90 ± 10 |

*Drosophila* ovaries consist of chains of egg chambers, each containing a developing oocyte and 15 germline nurse cells covered by a monolayer of somatic follicle cells. RNA FISH using a probe targeting the R2short sequence showed accumulation of R2short-rRNAs in *piwi* mutant nurse cells, as compared to control heterozygous sisters (Fig. 5B), but not in the developing oocyte, which becomes transcriptionally inactive from stage 5 of oogenesis.^58^ The preferential derepression of R2short-rRNA in nurse cells may be linked to their exceptionally high levels of rDNA transcription, which are required to sustain ribosome production for oocyte maturation.^44^ Consistent with our subcellular fractionation results, the R2short FISH signal was observed mostly within nucleoli, which in nurse cells form complex, thick network-like structures^44^ displaying reduced DAPI staining (Fig. 5C). The R2short colocalized with pre-rRNA but was restricted to distinct zones within the *piwi* mutant nucleolus (Fig. 5C), potentially corresponding to the locations of R2short-rDNA units.

Previously, we reported that some *piwi* mutants exhibit nucleolar defects in nurse cells.^59^ Specifically, these mutants show a reduced immunostaining signal for Udd in nucleoli, despite having normal total Udd levels. Udd signal intensity in the nucleolus has been linked to rRNA production levels and likely serves as an indicator of overall nucleolar transcription initiation activity.^55^ Here, we observed a more pronounced reduction in Udd signal, but not in fibrillarin, in nurse cells of *piwi* mutants carrying R2short insertions compared to those without them (Fig. 5D). Since piwi mutations affect Udd localization only in nurse cells, we quantified these effects using the germline-to-somatic Udd signal ratio within the same egg chamber (Fig. 5E). We found that in *piwi^2^*/*piwi^Nt^* mutants, the germline Udd signal was significantly lower than in ovaries of *piwi^3^*/*piwi^Nt^* mutants (p < 0.01, two-tailed Mann–Whitney U test). The *piwi^2^* and *piwi^3^* alleles originate from the same genetic background^54^, but only *piwi^2^* carries R2short-rDNA units on its X chromosome. In *piwi^Nt^*/*piwi^Nt^* combination which lacks R2short-rDNA units, but has different genetic background, we detected no significant effect of Piwi on Udd localization. These findings raise the possibility that the accumulation of aberrant rRNA molecules may impair nucleolar homeostasis.

### Piwi silences R2short-rDNA transcription

Using a nuclear run-on assay normalized to ovarian genomic DNA, we found that in control heterozygotes, the mean transcription rate per copy of intact rRNA genes without insertions was approximately two orders of magnitude higher than that of both long R2-rDNA and R2short-rDNA (Fig. 6A). This finding is consistent with previous reports that interrupted rDNA units are transcriptionally silenced.^21,22^ In *piwi^2^*/*piwi^Nt^* mutants, nascent 28S-R2short and 3′R2-28S junction RNAs increased ∼15-fold compared to control sisters (p < 0.01) (Fig. 6A), demonstrating that Piwi represses R2short-rDNA transcription. Although in control ovaries, the mean transcription rates per copy of R2short-rDNA and long R2-rDNA were similar (Fig. 6A), the steady-state nuclear level of R2short-rRNA per copy was more than seven times higher than that of long R2 (Fig. 6B). We then calculated nuclear RNA stability as the ratio of steady-state nuclear RNA to nascent RNA (Fig. 6C) and general RNA stability as the ratio of total steady-state ovarian RNA to nascent RNA (Fig. S8). Nuclear R2short-rRNA was ∼10 times more stable than long R2 RNAs, ∼100 times more stable than R1, and half as stable as intact nuclear 28S rRNA (Fig. 6C). These results suggest that the nuclear RNA surveillance machineries more effectively eliminate long R1 and R2 RNA, but are unable to recognize or degrade R2short-rRNA. Previously, one study highlighted that transcripts produced from the 167 bp 3′-terminal R2 fragment are more abundant than all other R2 RNAs,^15^ implying that similar R2short insertions in other lines also lead to the accumulation of long-lived transcripts. Notably, the stability of R2short-rRNA was unchanged in *piwi* mutants compared to control (Fig. 6C), supporting the idea that Piwi-piRNA complexes only reduce R2short-rRNAs transcription and not stability.

**Fig. 6.**
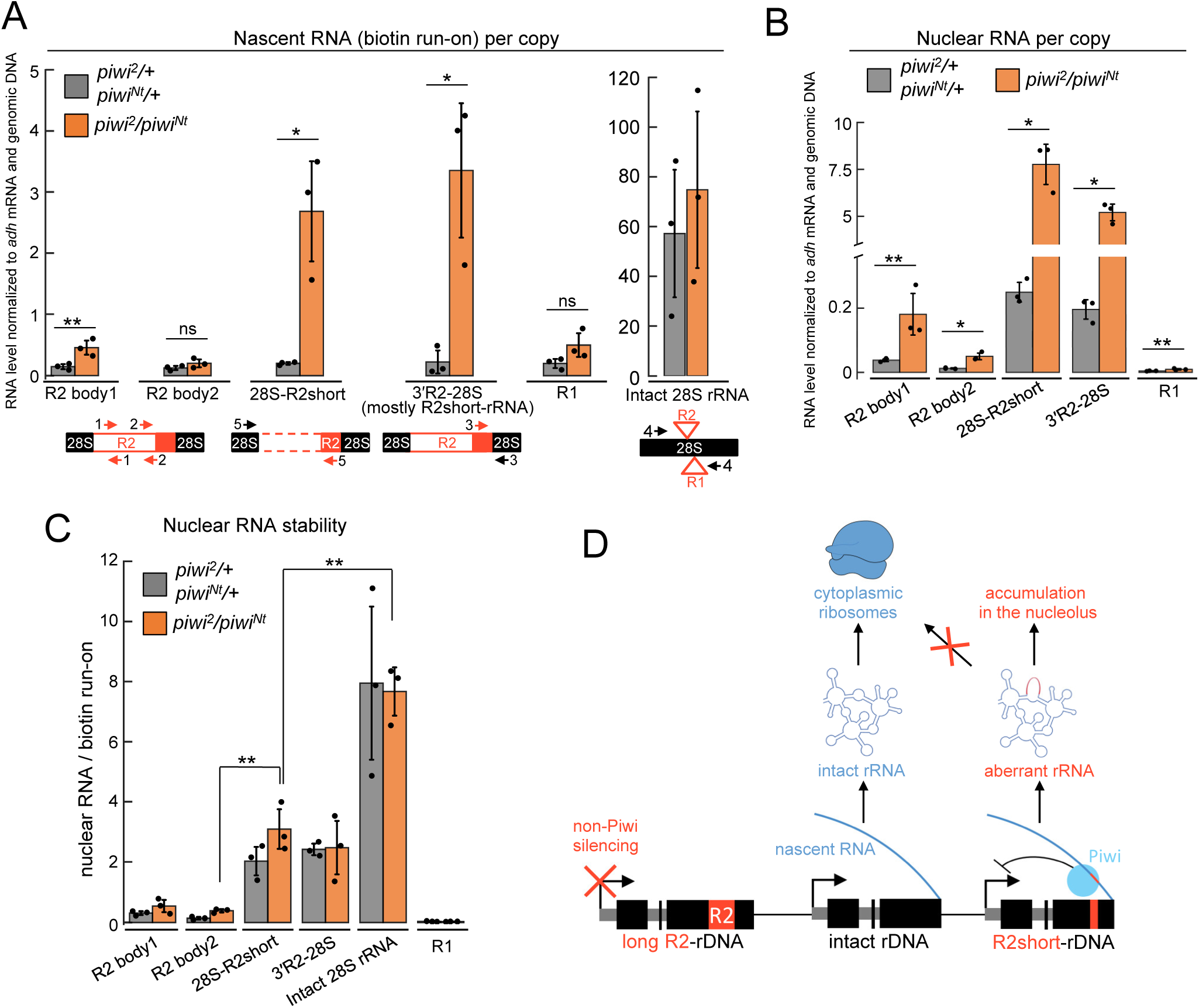
Piwi silences R2short-rDNA transcription. **A** Nascent RNA levels in ovaries of *piwi^2^/piwi^Nt^* and heterozygous sisters evaluated by biotin run-on, normalized to *adh* mRNA and genomic DNA. Numbers and positions of primers are shown. **B** Levels of steady-state nuclear RNA normalized to *adh* mRNA and genomic DNA. **C** Nuclear RNA stability calculated as the ratio of steady-state nuclear RNA to biotin run-on values. In all panels, p-value: *<0.01, **<0.05, two-tailed t-test, ns = not significant. **D** Working model of R2short-rRNA repression by Piwi.

## Discussion

### Piwi-dependent and independent silencing of interrupted rDNA units

In flies, many rDNA repeats carry insertions of full-length or truncated non-LTR retrotransposons R1 and R2. Transcription of rRNA genes containing TE fragments can generate broken and non-functional rRNA transcripts. Repressing such defective rDNA units is likely essential, especially in ovarian nurse cells and other tissues with a high demand for ribosome synthesis.

The piRNA-pathway represses TEs in the germline preventing transposition and ensuring genome stability.^32,33^ Here, we show that Piwi, loaded with antisense piRNAs, directs transcriptional silencing of aberrant rRNA in the nucleolus, preventing the production of defective pre-ribosomes. In *piwi* mutants, rDNA units containing 3′ fragments of the R2 element, R2short, are derepressed, leading to 28S rRNAs with embedded R2short sequences. The source of antisense R1 and R2 piRNAs is unclear. We propose that damaged rDNA fragments enriched with TE sequences, located in heterochromatic regions distinct from the rDNA cluster^60^, rather than interrupted rDNA units within the NOR, may generate these antisense piRNAs.

Piwi loss has little or no effect on the expression of long R1- and R2-rDNAs, even though these elements are complementary to abundant Piwi-bound antisense piRNAs. However, when overexpression of long R2-rRNAs and R1-rDNAs is triggered by depletion of the Pol I transcription factor, Piwi partially silences them. We propose that rDNA units with long insertions are simultaneously repressed by both Piwi-mediated and Piwi-independent mechanisms (Fig. 6D). The nature of other mechanisms for recognizing interrupted rDNA remains a mystery. Hypothetically, these mechanisms might detect misfolded or fragmented nascent rRNA, inducing transcriptional silencing of the corresponding rDNA unit. Due to its small size and the absence of the R2 ribozyme, R2short-rRNA may escape detection by these mechanisms.

Consistent with the current model for TE silencing by piRNAs, Piwi silences R2short-rDNA transcription but does not affect transcript stability. We do not yet know how Piwi represses R2short-rDNA transcription or whether this process shares similarities with Piwi-mediated silencing of conventional Pol II-driven TEs. Unlike TEs outside of the nucleolus,^37,42,43,46^ canonical heterochromatin marks likely play a limited role in rDNA silencing: loss of H3K9 methyltransferases or the heterochromatin-binding protein HP1 has little impact on Pol I-dependent transcription of R1 and R2.^22,30^ In contrast, depletion of SUMO and SUMO ligases results in a 2-3 order of magnitude derepression of R1 and R2 that is stronger that derepression of other TEs.^30,31^ Piwi has been linked to the SUMOylation pathway,^61–63^ raising the possibility that SUMOylation could play a key role in Piwi-dependent silencing of interrupted rDNA units.

### A potential role of the Piwi-piRNA pathway in quality control of rRNA

The quality of eukaryotic rRNA is known to be controlled by several posttranscriptional RNA surveillance pathways, including degradation of erroneous or misprocessed rRNAs by the nuclear exosome^64,65^ and cytoplasmic non-functional rRNA decay (NRD) due to translation failure^66,67^. Much less is known about the mechanisms that prevent the transcription of abnormal rDNA.

We found that R2short-rRNA is relatively stable in the nucleolus (Fig. 6C), suggesting that it cannot be efficiently removed by post-transcriptional RNA surveillance, likely due to its structural integrity. Most R2short-rRNA molecules are not incorporated into mature cytoplasmic ribosomes (Fig. 6D). However, our results propose that their accumulation in nucleolar pre-ribosomes could be detrimental to nucleolar homeostasis. Thus, Piwi-mediated repression of R2short-rDNA may be biologically significant and contribute to the ovarian phenotype observed in *piwi* mutants.

A variety of small RNAs mapping to rRNA in both sense and antisense orientations have been identified across eukaryotes. Some of these RNAs play roles in the establishment of nucleolar dominance in plants and the maintenance of rDNA copy number in fungi.^68^ Notably, siRNAs have been shown to repress production of aberrantly elongated 5S rRNAs in *Arabidopsis*.^69^ In *C. elegans*, erroneous rRNAs can be utilized by RNA-dependent RNA polymerases to synthesize antisense ribosomal siRNAs (risiRNAs), which then reduce rDNA transcription, though the purpose of this regulation remains unclear.^70^ Interestingly, the piRNA pathway in *C. elegans* prevents the amplification of risiRNA populations.^71^ Since the genomes of various organisms harbor numerous compromised rDNA units, including rDNA-derived pseudogenes, it would be intriguing to investigate how widespread small RNA-mediated silencing of these sequences is across different species and to what extent it contributes to ensuring the production of functional rRNA.

## Methods

### *Drosophila* strains, maintenance and crosses

*Drosophila melanogaster* stocks and crosses were maintained under standard conditions at 25°C. For all experiments ovaries were dissected in phosphate-buffered saline (PBS) from adult female flies at age 3–7 days. For the analysis of *piwi* trans-heterozygous mutants, we used *piwi^2^* and *piwi*^06843^ (*piwi^3^*) null mutations^54^ and *piwi^Nt^* mutation with disrupted Piwi nuclear localization^38^. *piwi^Nt^/piwi^Nt^* females were collected from a strain with a higher survival rate of homozygous flies due to a change of genetic background^72^ compared to originally described *piwi^Nt^* line^38^. Piwi (#101658) and Udd (#25313) UAS-RNAi stocks were obtained from Vienna Drosophila Resource Center (VDRC). Germline knockdowns (GKD) were induced by crossing these lines with nos-GAL4 driver *P{UAS- Dcr-2.D}1, w*^1118^*, P{GAL4-nos.NGT}40* (#25751) from Bloomington Drosophila Stock Center (BDSC). To obtain Udd Piwi double knockdowns, we first generated *UAS-Piwi- RNAi/CyO*; *UAS-Udd-RNAi/TM3, Sb*^1^*, Ser*^1^ flies and then crossed them to the nos-GAL4 driver. Four analyzed genotypes (control non-KD, *piwi* KD, *udd* KD, and both *piwi* and *udd* KD) were obtained as female siblings, ensuring they have the same rDNA content on their X chromosomes.

### Analysis of small RNAs

Small RNA datasets were retrieved from the NCBI GEO database. Accession numbers and corresponding references for all datasets analysed in this study are listed in Table S1. To analyse 3’-end protection of R1-, R2-, and rDNA-derived small RNAs, we used oxidized and non-oxidized small RNA libraries prepared from the same samples of wild-type ovaries.^73^ As other non-mutant ovarian total small RNA libraries, we utilized control germline knockdown ovaries^74^, the MD wild type line^48^, and *w*^1^ line^49^. For reanalysis of small RNA coimmunoprecipitating with PIWI proteins we used datasets obtained for MD wild type line^48^; *w*^1^ line^49^; and *w*^1118^ line^50^. Removal of adapter sequences, 2S rRNA and poly(A) tails as well as filtering by quality or size ranging from 18 to 29 nt was done using cutadapt (version 4.9). Processed reads were then mapped to *D. melanogaster* genome (dm6), rDNA consensus (Document S2),^75^ R1 (GenBank: X51968.1), and R2 (GenBank: X51967.1) sequences using Bowtie2 version 2.4.4 with default settings. Further filtering and indexing were carried out using SAMtools version 1.17 (http://www.htslib.org/). The mapped reads were then sorted and separated by strand to calculate coverage for the plus and minus strands with SAMtools depth. The coverage data was saved in separate files. Coverage plots, read length histograms and WebLogo plots were carried out in R. ggseqlogo and gridExtra were utilized for generating sequence logos. To estimate the abundance of germline-derived piRNAs, processed reads from the dataset with germline knockdowns^76,77^ were mapped to 158841 bp *flamenco* reference sequence using Bowtie2. *flamenco* gene sequence was retrieved from the *D. melanogaster genome* (dm6). Obtained read counts were set as a “per million” scaling factor and used for normalization RPM values for rDNA, R1, and R2 sequences.

### RT-qPCR

TRIzol reagent (Invitrogen) was used to extract total RNA from hand-dissected ovaries and nuclear RNA from the nuclear pellet (see Biotin run-on assay). At least three biological replicates of RNA were used. DNA was removed by DNA Removal Kit (Invitrogen, AM1906) according to the manufacturer’s instructions. Nucleic acid quantification was done using NanoDrop 1000 spectrophotometer V3.8 or using the Qubit RNA BR Assay Kit (Invitrogen). 1 μg of RNA was used for the reverse transcription reaction with random hexamer primers and Mint reverse transcriptase (Evrogen), adhering to the manufacturer’s protocol. The resulting cDNAs were analyzed in triplicates on DT96 real-time DNA amplifier (DNA-Technology) or MJ Mini thermal cycler (Bio-Rad) using SYTO™ 13 Green Fluorescent Nucleic Acid Stain (Invitrogen) and Hot Start Taq DNA polymerase (Evrogen) applying a program of 94°C, 5 min, followed by 40 cycles at 94°C, 15-30 s (depending on primers); 60°-64°C (depending on primers), 20 s; 72°C, 20- 40 s. A melting curve analysis was performed from 72°C to 92°C, with 0.3°C increments. Relative expression was calculated from Cq values using a ΔΔCq method. PCR products were verified by gel electrophoresis and by Sanger sequencing. For the design of primers, the following sequences were used: R2 (X51967.1), R1 (X51968.1), rDNA clone pDm238 (NCBI M21017.1), *D. melanogaster* rDNA consensus^75^, which was originally downloaded from http://www.rochester.edu/college/bio/thelab and is provided as a supplementary material (Document S2). PCR primers and amplicon lengths are shown in Table S3.

Strand-specific reverse transcription was performed using 0.25 pmol of gene-specific primers and Superscript II reverse transcriptase (Invitrogen) according to manufacturer’s instructions. Control reactions were conducted without adding an RT primer.

### R1 and R2 isoform analysis in genomic DNA

The presence of truncated R1 and R2 insertions in analyzed *Drosophila* lines was examined by long-extension PCR of genomic DNA^17^ using primers flanking R1/R2 insertion sites in 28S rDNA sequence (pair #4, Table S3). Genomic DNA was isolated from whole flies according to the standard procedure.^78^ PCR reaction was conducted using 0.05 u/μl HotTaq Pol (Sileks) on 13 ng/μl genomic DNA, applying a program of 94°C, 10 min, followed by 30 cycles of amplification, each consisting of: 94°C, 40 s; 60° 1 min; 72°C 10 min. Under these conditions, the rate of the PCR reaction resembles a pseudo-first-order reaction, causing the amount of product to change with the concentration of the input DNA without preferential accumulation of short amplicons. PCR products were then run on an agarose gel, and all individual PCR bands up to 1000 bp were purified from the gel and Sanger sequenced from both ends. Sequencing confirmed that the lower PCR product contained intact 28S rDNA without insertions, while all upper bands contained R1 and R2 3′ fragments inserted in typical positions within the 28S rDNA. Additionally, genomic DNA was analyzed by long-extension PCR using a forward primer located in the upstream 28S rRNA sequence and a reverse primer in the 3′ end of R2 (pair #5, Table S3) to detect only R2 insertions in 28S rRNA. Sanger sequencing showed that the shortest band detected by pair #5 primers in *piwi^2^*-derived lines had a sequence identical to R2short detected by primer pair #4. This sequence was also identical to the single amplicon produced by primer pair #4 in RT-PCR under standard conditions (short PCR extension time) using ovarian cDNA from *piwi^2^*-derived lines.

### Biotin run-on assay

About 200 ovaries of *piwi^2^/piwi^Nt^* and their heterozygous sisters (*piwi^2^*/+, *piwi^Nt^*/+) were homogenized in 0.5 ml of homogenization buffer (10 mM Tris-HCl, pH 7.5, 300 mM sucrose, 3 mM MgCl2, 3 mM CaCl2, 10 mM NaCl, 0.1% (v/v) Triton X-100, 0.1% (v/v) NP40, 0.5 mM DTT, and 1× cOmplete Protease Inhibitor Cocktail (Roche)) using Dounce homogenizer (15 strokes with pestle B). Undestroyed tissues were removed by filtration through Miracloth (Merck Millipore). Nuclei were pelleted by centrifugation for 5 min at 1000×g at 4°C, washed with homogenization buffer, and resuspended in 240 µl of resuspension buffer (Tris-HCl, pH 8.0, 25% glycerol, 5 mM MgCl2, 0.1 mM EDTA, 5 mM DTT, and 1× protease inhibitor cocktail). 20 µl of the nuclei suspension (nuclear input sample) was used for subsequent RNA isolation using the TRIzol-LS (Invitrogen) according to the manufacturer′s protocol and RT-qPCR analysis. 100 µl of the nuclei suspension were mixed with 100 µl of biotin+ reaction solution (10 mM Tris-HCl, pH 8.0, 5 mM MgCl2, 0.1 mM DTT, 0.3 M KCl, 0.25 mM ATP, 0.25 mM GTP, 0.05 mM Biotin-11-UTP (PerkinElmer), 0.05 mM Biotin-11-CTP (PerkinElmer), 80 U RiboLock RNase Inhibitor (Thermo Fisher Scientific)). As a biotin- control, 100 µl of the nuclei suspension were mixed with 100 µl of the same solution containing UTP and CTP instead of biotinylated analogs. The reaction proceeded for 7 min at 30°C and was stopped by adding the TRIzol-LS, with subsequent RNA isolation according to the manufacturer′s instructions. Biotin-labeled newly synthesized RNA was precipitated using Dynabeads kilobaseBINDER kit (Thermo Fisher Scientific) according to the manufacturer′s instructions. After washings, RNA-bound beads were resuspended in water. Then cDNA synthesis was conducted directly on beads with random hexamer primers and Mint reverse transcriptase (Evrogen), according to the manufacturer’s protocol. The synthesized cDNA was released by heating to 95°C for 10 min and used for qPCR analysis. Negative bio- controls were analyzed by qPCR for all primer pairs in parallel for all replicas. The absolute qPCR values for the bio- controls were at least 10 times less than for bio+ samples. To evaluate average per-copy transcription rates or per-copy steady-state RNA level for various types of rDNA units, R1 and R2 TEs, we isolated genomic DNA from at least from 3 biological replicas of ovaries according to the standard procedure.^78^ qPCR of genomic DNA was conducted using the same primers and conditions as RT-qPCR and normalized to *rp49* genomic DNA.

### Evaluation of relative per-copy transcription rates, steady-state RNA levels, and RNA stability

To assess the average per-copy transcription rates and steady-state RNA levels of various rDNA units, as well as R1 and R2 transposable elements (TEs), we performed qPCR on genomic DNA from at least three biological replicates using the same primers and conditions as RT-qPCR, normalizing to *rp49* genomic DNA. The average genomic qPCR values were calculated for each primer pair. Run-on or RT-PCR values were then normalized to genomic qPCR values for the corresponding sequence and genotype. Run-on or RT-qPCR values for the 3′R2-28S junctions (pair #3, Table S3) were normalized to 28S-R2short genomic DNA PCR values (pair #5, Table S3), as 3′R2-28S cotranscripts are predominantly produced by R2short insertions, while all R2 insertions contain the same 3′R2-28S junctions in DNA.

Nuclear RNA stability was estimated as the ratio of RT-qPCR values from nuclear input samples to those from biotin+ nascent RNA samples within the same biological replicate. General RNA stability was calculated as the ratio of total steady-state ovarian RNA to nascent RNA.

### Isolation of ribosomes and pre-ribosomes

For isolation of cytoplasmic mature ribosomes, 50–100 ovaries of *piwi^2^/piwi^Nt^* mutants were homogenized in 1.4 ml of homogenization buffer (20 mM HEPES -NaOH, pH 7.6, 100 mM KCl, 10 mM MgCl2, 0.5% (w/v) sodium deoxycholate, 1% (v/v) NP40, 0.1 mg/ml cycloheximide, 1 mM DTT, 1:200 RiboLock RNase Inhibitor, 1× protease inhibitor cocktail) using Dounce homogenizer (30 strokes with pestle A and the same with pestle B). 100 µl of the homogenate was used as an input sample for RNA isolation using the TRIzol-LS (Invitrogen). The rest of the homogenate was centrifuged at 16,000 × g for 20 min at 4°C to remove undestroyed tissues, nuclei, and other large cellular components. The supernatant was centrifuged at 100,000 × g for 2 h using Himac CP100NX ultracentrifuge, rotor P50A3-0529. Ribosome pellet and 300 µl of supernatant after ultracentrifugation (designated as Cytosol) were used for RNA isolation using the TRIzol-LS reagent (Invitrogen). After isolation, RNA samples were treated with DNase using the DNA-free DNA Removal Kit (Invitrogen) and used for cDNA synthesis and qPCR analysis as described above in RT-qPCR section. 0.5 µg of RNA from each fraction (Input, Cytosol, Ribosomes) was used for reverse transcription.

Nuclear pre-ribosomal complexes were purified according to the published protocol^79^ with modifications. 200–250 ovaries were homogenized in 0.5 ml of buffer, containing 10 mM Tris-HCl, pH 7.5, 300 mM sucrose, 10 mM MgCl2, 100 mM KCl, 3 mM CaCl2, 10 mM NaCl, 0.1 mg/ml cycloheximide, 0.1% (v/v) Triton-X100, 0.1% (v/v) NP40, 1 mM DTT, and 1× protease inhibitor cocktail using Dounce homogenizer (15 strokes with pestle B). Undestroyed tissues were removed by centrifugation at 1000 × g through Miracloth membrane (Merck Millipore). Then nuclei were pelleted by centrifugation for 5 min at 1000 × g at 4°C and washed with homogenization buffer. The nuclear pellet was resuspended in the above-described homogenization buffer for ribosome extraction and homogenized rigorously using a Dounce homogenizer. The nuclear lysate was then centrifuged at 21,000 × g for 1 h to sediment large nuclear fragments including chromatin. The pellet was discarded and the supernatant was ultracentrifuged at 100,000 × g for 2 h, resulting in a pellet enriched for pre-ribosomes and a supernatant containing the nucleoplasm. RNA was extracted from all fractions, and equal amounts were used for RT-qPCR.

To calculate the relative abundance of transcripts in subcellular fractions, we used equal amounts of RNA for RT-qPCR from total ovaries (Input), cytosol, cytoplasmic ribosomes, and nuclear RNA, which was isolated in a separate experiment. RT-qPCR values for cytosolic, ribosomal, and nuclear RNA were normalized to the Input RT-qPCR value for the corresponding experimental replicate. Alternatively, taking into account the total RNA yield from each fraction, we calculated the percentage of molecules in each fraction by setting the Input as 100%. To determine the percentage of molecules in nuclear pre-ribosomes and the nuclear supernatant, we estimated the total number of molecules based on RT-qPCR for each fraction, considering the fraction of cleared nuclear lysate (i.e., the nuclear extract after removal of large nuclear fragments) as 100%.

### Nanopore sequencing and data treatment

High molecular weight DNA (HMW DNA) was extracted from ∼100 ovaries of *piwi^2^* line with the genotype *w*^1118^*; P{ry11}piwi*^2^*/CyO* using Monarch HMW DNA Extraction Kit for Tissue (NEB). DNA concentration was measured using a Qubit fluorimeter after shearing according to ONT Ultra-Long DNA Sequencing Kit protocol (Oxford Nanopore Technologies). The sequencing library was prepared from 10 μg of HMW DNA using Ultra-Long DNA Sequencing Kit (SQK-ULK001, Oxford Nanopore Technologies) according to manufacturer protocol (Oxford Nanopore Technologies, version: ULK_9124_v110_revE). Volumes were scaled down 3× to match the gDNA amount. Flow cell priming and library loading were performed according to SQK-ULK001 protocol. Sequencing run lasts for 48 h with 6h mux scan intervals. The library was divided into two parts (5 µg each) and loaded twice with DNase wash in between using Flow Cell Wash Kit EXP-WSH004 kit. A MinION R 9.4.1. flow cell (Oxford Nanopore Technologies) was used to sequence the library without base calling. Sequencing of *w*^1118^*; P{ry11}piwi*^2^*/P{ry11}piwi*^2^ whole flies was performed using a Ligation Sequencing Kit (Oxford Nanopore Technologies SQK-LSK109) as described.^80^

Fast5 files generated by sequencing were base called using the dna_r9.4.1_450bps_hac profile in Guppy (version 3.5.2) running on a standalone GPU-enabled server. Reads were filtered by length using NanoFilt, and reads ≥ 10,000 bp were used for subsequent analysis. Reads were then blasted using BLASTn against rDNA elements (18S, 5.8S, 28S) retrieved from *the D. melanogaster* rDNA consensus sequence^75^. Matched reads were extracted to a separate library. IGS 96bp, 240bp, and 330bp repeats (retrieved from rDNA clone pDm238 (NCBI M21017.1)), R1 (GenBank: X51968.1), R2 (GenBank: X51967.1), and other TE sequences (retrieved from transposon_sequence_set_flybase_072020.fa) were mapped to the subset of ONT reads containing rRNA sequences by BLASTn and converted to bed files using blast2bed. Regular expressions were utilized to sort nanopore reads into directories depending on which TE they contained. Nanopore reads with features representing various elements from the bed files were visualized in the IGV browser (version 2.12.3). To find R2 insertions, all reads containing R2 sequences were extracted and visually inspected in the IGV browser.

### RNA FISH and Immunostaining

RNA FISH with probe to R2 and pre-rRNA was carried out as previously described^22^ using Cy3- and Cy5-labeled oligonucleotide probes, respectively (Table S3). Immunostaining was done as described^81^ immediately after fixation. The following primary antibodies were used: rabbit anti-fibrillarin (1:1000, Abcam #5821); guinea pig anti-Udd (1:800, provided by M. Buszczak)^55^. Anti-rabbit IgG Alexa Fluor 546 and anti-guinea pig IgG Alexa Fluor 488 (Thermo Fisher Scientific) were used as secondary antibodies. Imaging was conducted by LSM 900 Confocal with Airyscan 2 super-resolution detector (Zeiss) was used for. Images were processed and analyzed using IMARIS 7.4.2 software (Bitplane AG).

### Quantification of confocal signal intensity

To quantify Udd signal in control and mutant ovaries, grayscale confocal images acquired under identical settings were analyzed using Fiji (version 2.16.0/1.54g). For each genotype, we analyzed 13-17 egg chambers of stages 4-5, measuring Udd nuclear signals in 4–7 nurse cells and 7–10 somatic follicle cells per egg chamber. Nuclear areas were selected using polygon selection based on the DAPI signal. Measurements included area, mean gray value, and integrated density for each nucleus and the image background. Corrected mean gray values were calculated using the following formula: Corrected mean gray value of nucleus = (integrated density of nucleus – mean gray value of background × area of nucleus)/area of nucleus. For each egg chamber, Udd signal values for nurse cells and somatic follicular cells were averaged, and the germline-to-somatic Udd signal ratio was calculated.

### Statistic and reproducibility

All data are presented as the mean ± standard deviation (SD), unless indicated otherwise. The exact number of biological replicates (n) is provided in the figures. Statistical tests used for each figure are indicated in the corresponding figure legends; unless otherwise specified, comparisons were performed using a two-tailed unpaired Student’s t-test, with p < 0.05 considered statistically significant. Differences in confocal signal intensity were assessed using a two-tailed Mann–Whitney U test. Statistical analyses were performed using Microsoft Excel (Office 16) or GraphPad Prism 8 (GraphPad Software, San Diego, CA, USA). No statistical methods were used to predetermine sample size. Investigators were not blinded during experiments or outcome assessment.

## Supporting information

Sapplementary figures

## Data availability

All data and materials are available from the corresponding authors upon request. The authors declare that all data supporting the findings of this study are available within the article and Supplementary Information files. The Nanopore sequencing datasets are available from the National Center for Biotechnology Information Sequence Read Archive under the accession number PRJNA1114025.

## Acknowledgements

We thank Vladimir Gvozdev, Anastasia Stolyarenko, and members of the Zamore laboratory for help and discussions, Michael Buszczak for Udd antibodies. This work was supported by the National Institutes of Health grant R35 GM136275 to P.D.Z. and by the Russian Science Foundation (RSF) grant 19-14-00382 to M.S.K. The work of A.S.S. was conducted under the IDB RAS Government basic research program in 2025 № 0088-2024-0017.

## Author contributions

Conceptualization: M.S.K. and E.A.F. Methodology: M.S.K., A.S.S., S.A.P., and P.D.Z. Investigation: E.A.F., A.S.S., E.A.M., Y.A.A., S.A.L., S.A.P., V.A.P., A.A.I., and M.S.K. Supervision: M.S.K. Funding acquisition: M.S.K. and P.D.Z. Writing – original draft: M.S.K. Figure preparation: M.S.K. and E.A.F. Writing – review and editing: M.S.K. and P.D.Z. with input from all co-authors.

## Competing interests

The authors declare no competing interests.

**Correspondence** and requests for materials should be addressed to Mikhail S. Klenov.

