## Supplementary material for "Piwi and piRNAs repress transcription of aberrant rRNA genes containing retrotransposon fragments": Sapplementary figures

### Supplementary Figures 1-8 and Table S2

**for**

### Piwi and piRNAs repress transcription of aberrant rRNA genes containing retrotransposon fragments

Elena A. Fefelova[1,7], Aleksei S. Shatskikh[2,7], Elena A. Mikhaleva[1], Yuri A. Abramov[1], Sergey A. Lavrov[1], Sergei A. Pirogov[3], Valentin A. Poltorachenko[4], Artem A. Ilin[3], Phillip D. Zamore[5,6], Mikhail S. Klenov[1,5,*]

[1]Department of Molecular Genetics of the Cell, Institute of Molecular Genetics, Russian Academy of Sciences, Moscow 123182, Russia
[2]Koltzov Institute of Developmental Biology, Russian Academy of Sciences, 26 Vavilov Street, Moscow 119334, Russia
[3]Department of Molecular Biosciences, Stockholm University, The Wenner-Gren Institute, Stockholm, Sweden
[4]Department of Microbiology, Tumor and Cell Biology, Karolinska Institutet, Stockholm, Sweden
[5]RNA Therapeutics Institute, University of Massachusetts Chan Medical School, 368 Plantation Street, Worcester, MA 01605, USA
[6]Howard Hughes Medical Institute, University of Massachusetts Chan Medical School, 368 Plantation Street, Worcester, MA 01605, USA
[7]These authors contributed equally to this work

#### Figure S1

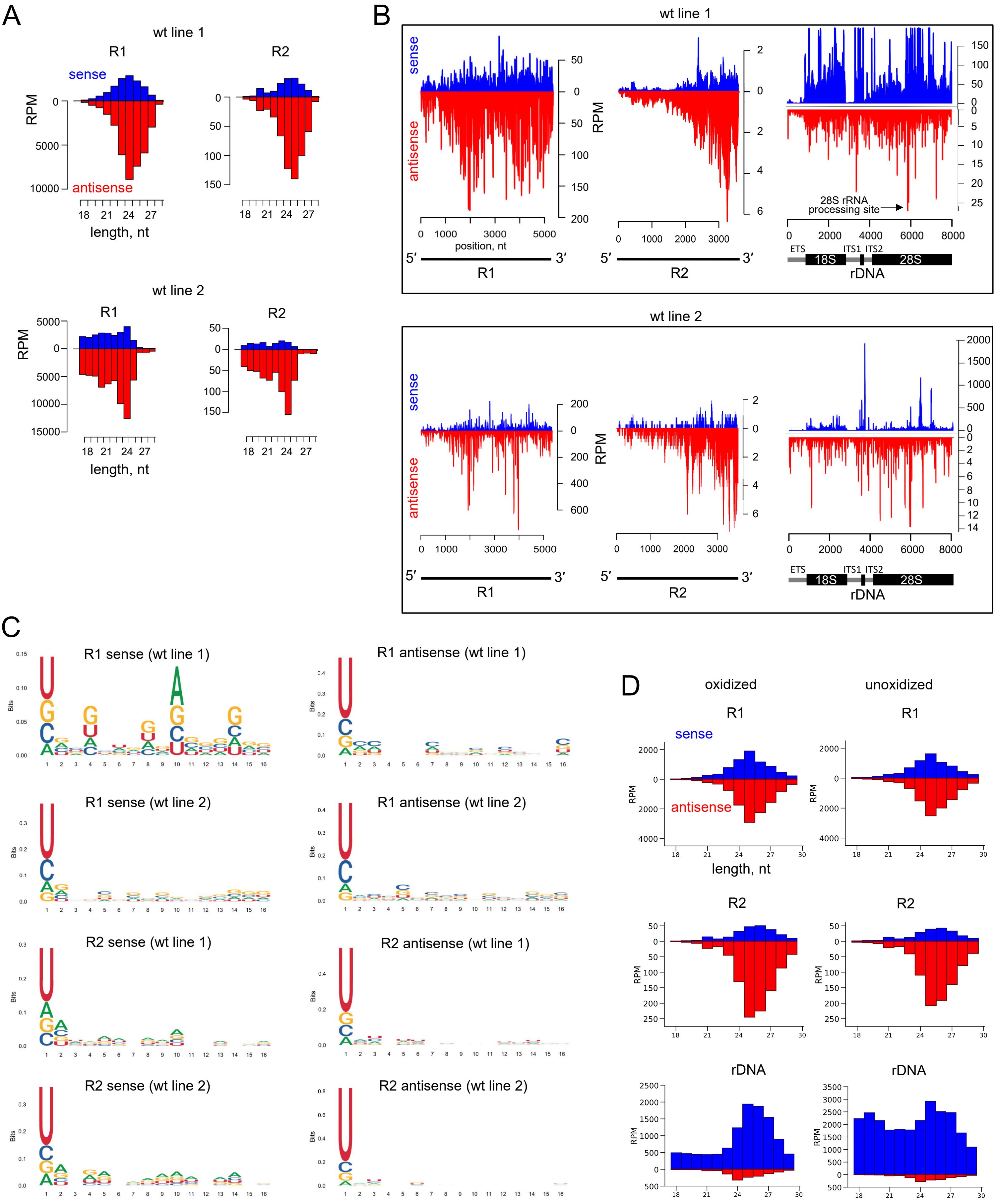


**Figure S1. Most ovarian R1 and R2 small RNAs are piRNAs.**
(**A**) Size distribution of ovarian small RNAs (18–29 nt) mapping to R1 and R2 transposable elements in control germline knockdown ovaries[74] (designated as wt line 1) and the MD wild type line[48] (designated as wt line 2). (**B**) Coverage of R1, R2, and rDNA sequences by ovarian small RNAs. Reads corresponding to 2S rRNA were removed. The highest peak of antisense rDNA reads corresponds to the site of 28S rRNA cleavage into 28Sa and 28Sb parts, a *Drosophila*-specific cytoplasmic rRNA processing step. (**C**) Logo plots of sense and antisense R1 and R2 small RNAs in ovaries of wt line 1 and wt line 2. (**D**) Comparison of R1, R2, and rDNA small RNA amount and size distribution in oxidized and non-oxidized libraries obtained from the same samples of wild-type ovaries[73].

### Figure S2

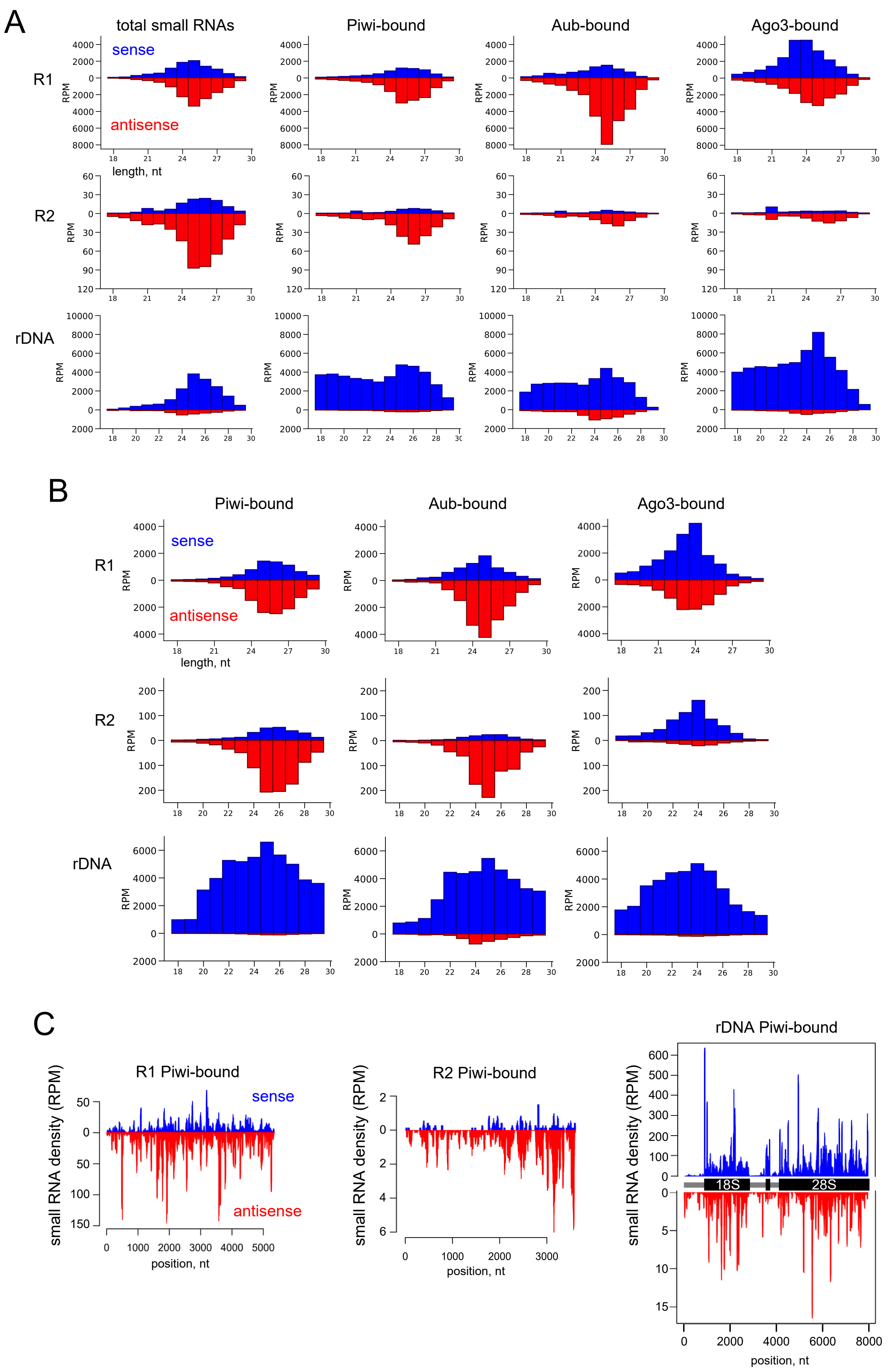


**Figure S2. R2 piRNAs are predominantly bound to Piwi.**
(**A**) Amount and size distribution of small RNAs corresponding to R1, R2 and rDNA isolated from immunoprecipitated Piwi, Aub, and Ago-3 proteins (reanalysis[49]). (**B**) Small RNAs corresponding to R1, R2 and rDNA isolated from immunoprecipitated Piwi, Aub, and Ago-3 proteins (reanalysis[50]). (**C**) Coverage of R1, R2, and rDNA sequences by small RNAs associated with immunoprecipitated Piwi (reanalysis[48]). Reads corresponding to 2S rRNA were removed.

### Figure S3

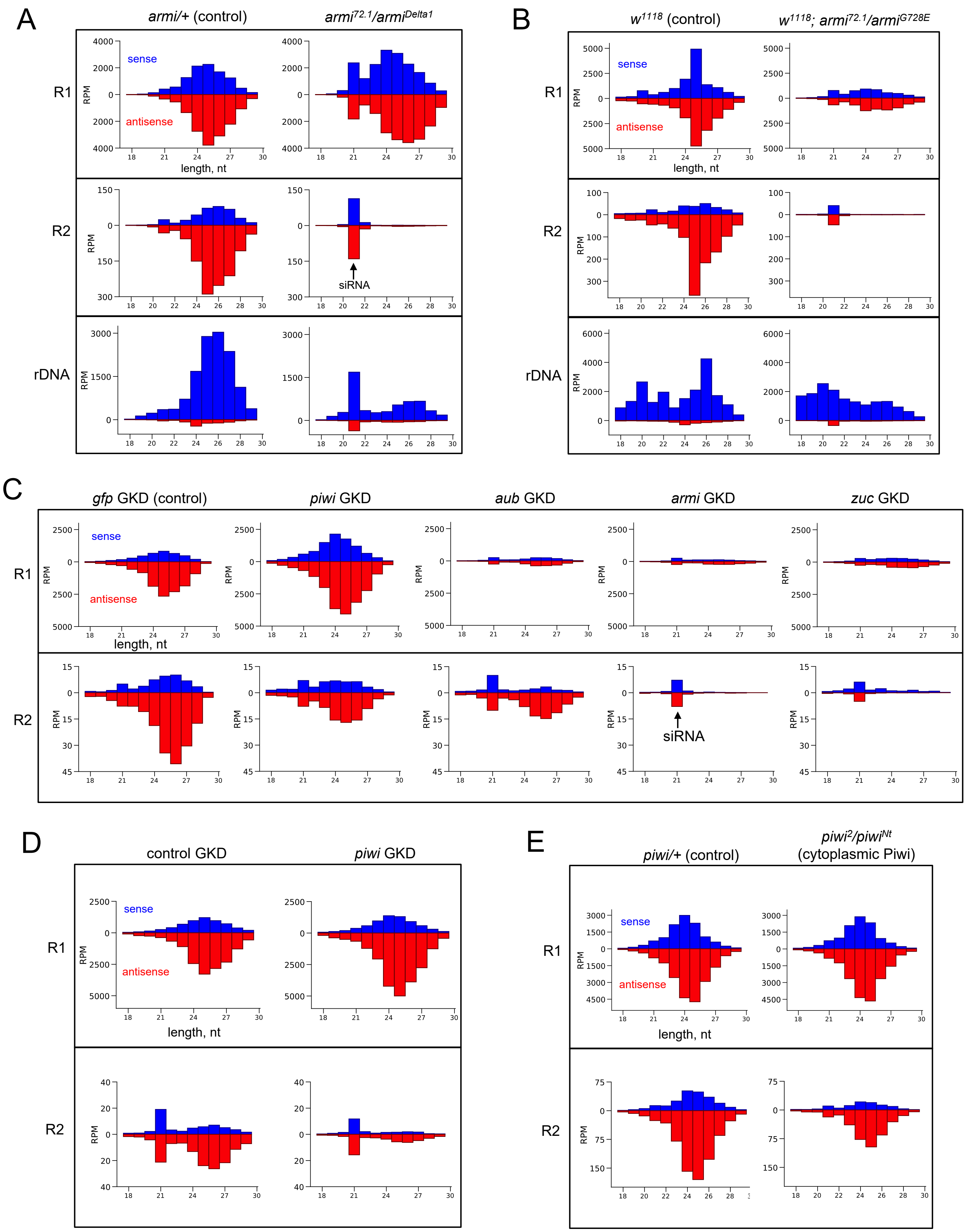


**Figure S3. Production of R2 piRNAs depend on Armitage (Armi) and occurs mostly in germ cells.**
(**A**) Amount and size distribution of small RNAs aligned to R1, R2, and rDNA in ovaries of *armi*$^{72.1}$/*armi*$^{Delta1}$ mutants and control heterozygotes (*armi*/+).[52] (**B**) Small RNAs in *w*$^{1118}$; *armi*$^{72.1}$/*armi*$^{G728E}$ mutants and control *w*$^{1118}$ ovaries.[73] (**C**) Small RNAs following MTD-GAL4-driven GKD of *piwi*, *aub*, *armi*, *zuc,* and control *gfp* GKD.[76] (**D**) Small RNAs following MTD-GAL4-driven germline knockdown (GKD) of *piwi*.[77] (**E**) Small RNAs in *piwi*$^{2}$/*piwi*$^{Nt}$ mutants and control heterozygotes.[73]

### Figure S4

A

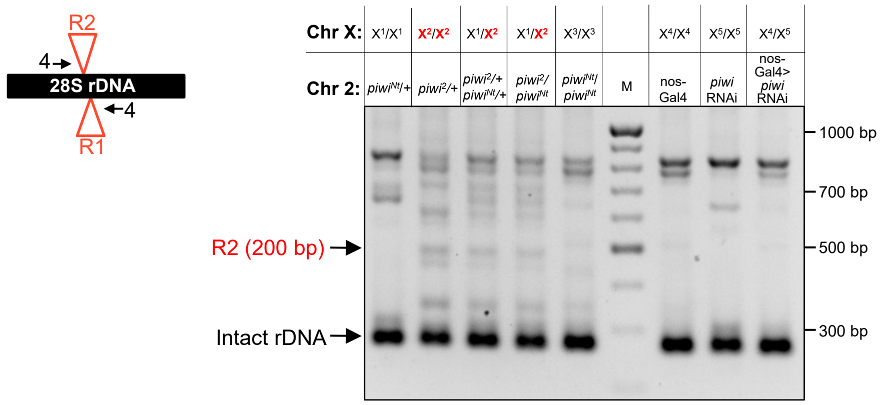


B

duplicated R2 sequence

upstream 28S rRNA

non-templated nt

R2 sequence

GAAATTCAAGTAAGCGCGGGTCAACGGTTATGGTTATAATTTTTTTTTTTTAATGGTTATAATTAAATGGCTATGAGGATGGTTTT

AGTACGTAGGCGTTGCGGAACTTCGGTTCATATAGAGCAATGAATCGTGCATGCTAGGAAAACTGACCACACACAGTGTTGGCA

downstream 28S rRNA

AACCTAGTATCTTTCGAAAATTTCCATACCCCCGCGATCAAAAAAAAAAAAAAAAAATAGCCAAATGCCTCGTCATCTAATTAGT

GACGCGCATGAATGGATTAACGAGATTCCTACTGTCCCTATCTACTATCTAGCGAAACCACAGCCAAGGGAACGGGCTTGGAAT

AATTAGCGGGGAAAGAAGACCCTTTTGAGCTTGACTCTAATCTGGCAGTGTAAGGAGACATAAGAGGTGTAGAATAAGTGG

**Figure S4. Detection of R2short insertions by long-extension PCR.**
(**A**) Location of primers #4 (black arrows) for analysis of R1 and R2 insertion length by long-extension PCR of genomic DNA and a gel image showing PCR products. The bottom band on the gel (268 bp) corresponds to intact 28S rDNA. For rDNA units containing insertions, the size of the PCR product increases by the length of the insertion. Genomes of the *piwi*$^2$ line and descendants carrying the X chromosome from the *piwi*$^2$ line (designated as $X^2$) contain R2short insertions of ~ 200 bp, which according to Sanger sequencing correspond to the R2 3′ fragment. Lines containing X chromosomes of other origin ($X^1$, $X^3$, $X^4$, $X^5$) lack R2short insertions. Other bands detected in such a range of sizes according to Sanger sequencing correspond to 3′ fragments of the R1 element. M: 1kb DNA ladder. (**B**) Sequence of the 28S rDNA fragment containing the R2short insertion, determined by Sanger sequencing of the genomic PCR band from the *piwi*$^2$ line. R2short is inserted at a typical R2 integration site and consists of 177 bp of the 3' UTR, preceded by an 11 bp non-templated poly-T tract and a 12 bp target site duplication.

### Figure S5

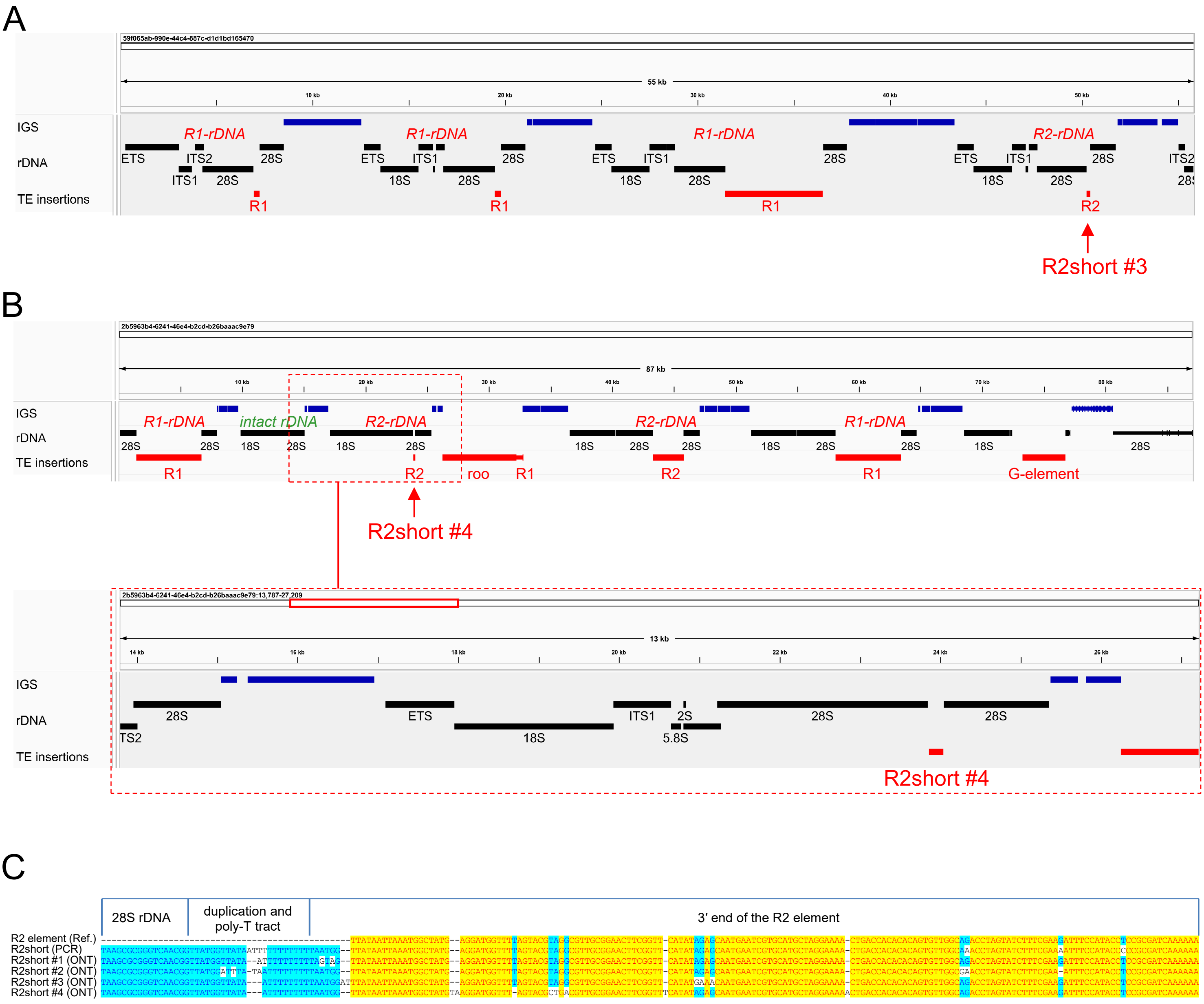


**Figure S5. Characterization of R2short-rDNA units by nanopore sequencing.**
(**A**) IGV browser screenshot showing the location of rDNA unit with R2short #3 insertion in 55 kb-long nanopore reads corresponding to fragments of the rDNA cluster in the *piwi*$^2$ line. Positions of intergenic spacers (IGS), rDNA, and TE insertions, including R1 and R2, are indicated. (**B**) Top: location of an rDNA unit with R2short #4 insertion in 87 kb-long nanopore read derived from the *piwi*$^2$ line rDNA cluster. Bottom: enlarged view of the rDNA-R2short #4 unit. (**C**) Alignment of R2short insertions extracted from nanopore reads (ONT), the R2short sequence obtained by Sanger sequencing of genomic DNA (PCR), and the reference sequence of the R2 3′ fragment (Ref.).

#### Figure S6

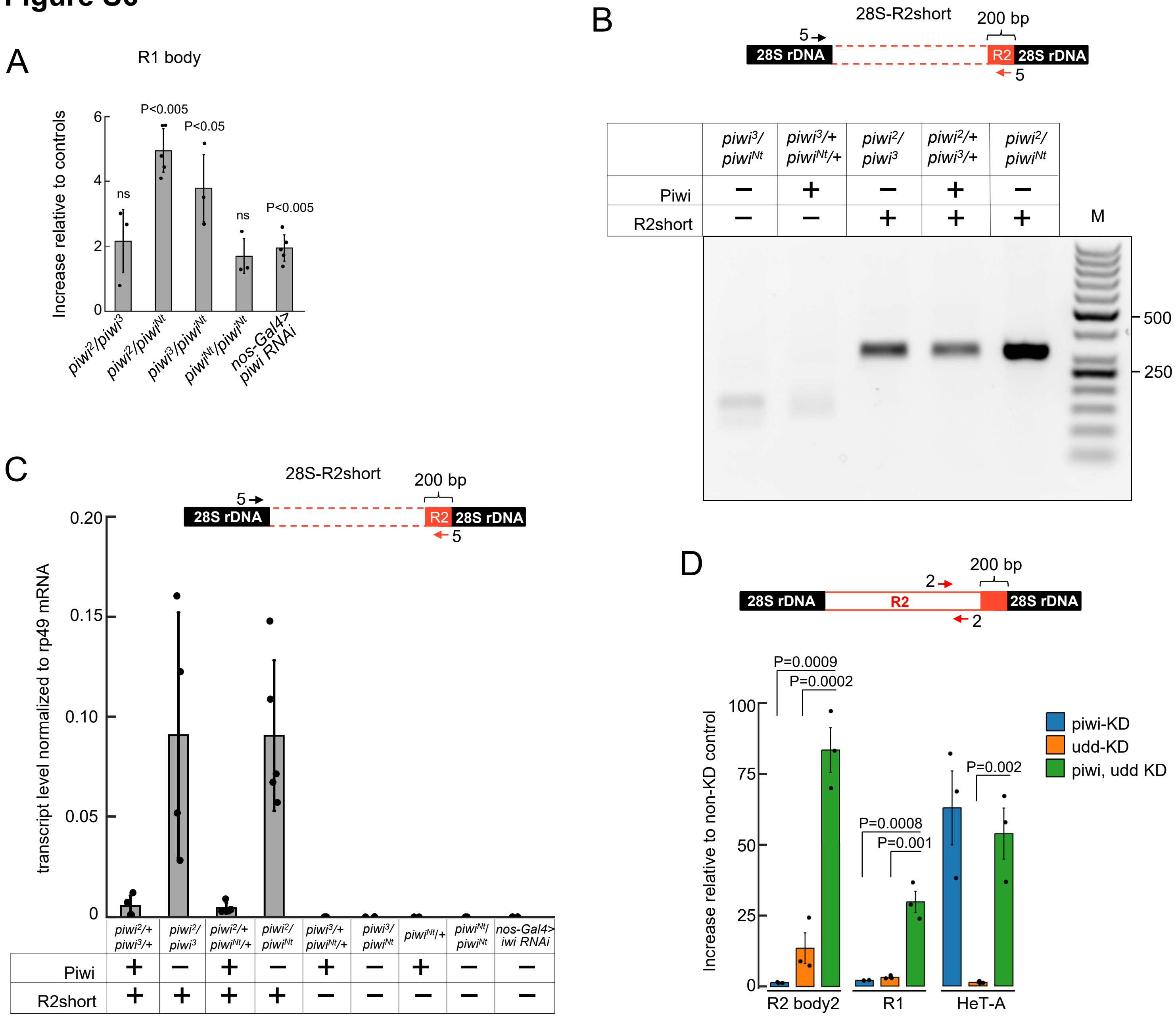


**Figure S6. Effect of the Piwi loss on the expression of R1 element and rDNA units carrying R2short and long R2 insertions.**
(**A**) Increase (fold change) of R1 element expression in *piwi* mutant combinations compared to heterozygous sisters carrying the same X chromosomes, normalized to rp49 mRNA. p-values are from a two-tailed t-test; ns = not significant. (**B**) Gel image of RT-PCR products corresponding to 28S-R2short transcripts (primers #5) in *piwi* mutant combinations. The presence (+) or absence (-) of R2short insertions is indicated based on genomic PCR analysis. The 28S-R2short product is detected only in lines with R2short insertions. (**C**) Abundance of 28S-R2short transcripts in different *piwi* mutant lines, normalized to *rp49* mRNA. In lines without R2short insertions, only background PCR levels are detected. (**D**) Increase of R2 body (primers #2), R1, and retrotransposon HeT-A expression in ovaries of nos-GAL4-driven germline knockdowns (KD) of *piwi*, *udd*, and both *piwi* and *udd* relative to control non-KD sisters. All genotypes carried the same X chromosomes with rDNA arrays. Expression of HeT-A, which is known to be silenced by Piwi, is not affected by *udd* KD. p-values are from a two-tailed t-test.

### Figure S7

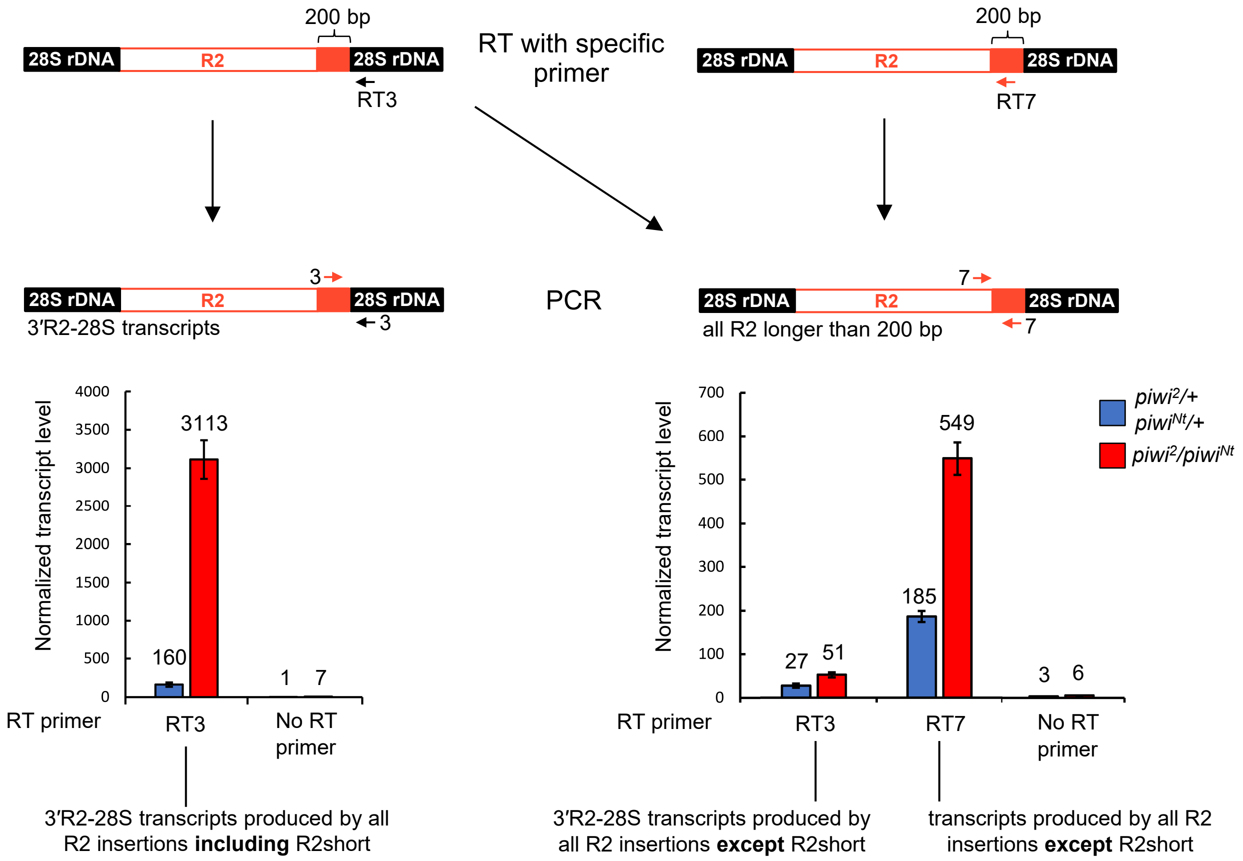


**Figure S7. R2short-rDNA units produce transcripts in which the R2 3′ end is joined to the downstream 28S rRNA sequence (3′R2-28S transcripts).**
Experimental setup and results of primer-specific RT-qPCR in *piwi*$^{2}$/*piwi*$^{Nt}$ and *piwi*$^{2}$/+ ovaries. PCR with primers #3 (left graph) detects 3′R2-28S transcripts produced by all R2 insertions, including R2short RNA, when using cDNA synthesized with the RT3 primer annealed to downstream 28S rRNA. PCR with primers #7 (right graph) on cDNA synthesized with RT3 primer detects 3′R2-28S transcripts derived exclusively from long R2 insertions. PCR with primers #7 using cDNA synthesized with RT7 primer detects all transcripts produced by long R2 isoforms. 'No RT primer' samples serve as a negative control, showing background levels of reverse transcription in the absence of a primer.

### Figure S8

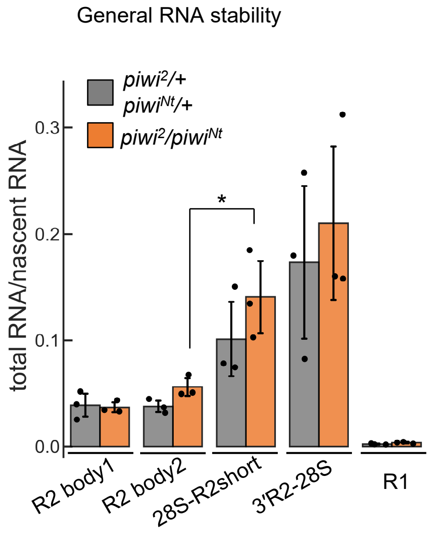


**Figure S8. General RNA stability of R2, R2-rRNA and R1 transcripts.**
General RNA stability was calculated as the ratio of ovarian steady-state RNA to nascent RNA in *piwi*$^{2}$/*piwi*$^{Nt}$ mutant ovaries and control heterozygotes. p-value: *<0.05, two-tailed t-test.

### Table S2

| | Total molecules as fraction of Input (%) | | | |
|---|---|---|---|---|
| | **Input** | **Cytoplasmic supernatant** | **Cytoplasmic ribosomes** | **Nuclei** |
| rp49 mRNA | 100 | 40 ± 20 | 14 ± 7 | 5 ± 1 |
| pre-rRNA (ITS1-5.8S) | 100 | 0.004 ± 0.002 | 1 ± 0.6 | 67 ± 8 |
| intact 28S rRNA | 100 | 0.002 ± 0.001 | 20 ± 10 | 15 ± 4 |
| 28S-R2short | 100 | 0.004 ± 0.004 | 2 ± 2 | 90 ± 30 |
| 3'R2-28S (mostly R2short-rRNA) | 100 | 0.007 ± 0.007 | 3 ± 1 | 100 ± 30 |
| long R2 (R2 body2) | 100 | 1.9 ± 0.5 | 2 ± 1 | 20 ± 10 |

**Table S2.** Total molecules as fraction of Input (%) in the cytosol, cytoplasmic ribosomes, and nuclei of $piwi^{2}/piwi^{Nt}$ ovaries.
